# Topographic Signatures of Global Object Perception in Human Visual Cortex

**DOI:** 10.1101/864736

**Authors:** Susanne Stoll, Nonie J. Finlayson, D. Samuel Schwarzkopf

## Abstract

Our visual system readily groups dynamic fragmented input into global objects. How the brain represents such perceptual grouping remains however unclear. To address this question, we recorded brain responses using functional magnetic resonance imaging whilst observers perceived a dynamic bistable stimulus that could either be perceived *globally* (i.e., as a grouped and coherently moving shape) or *locally* (i.e., as ungrouped and incoherently moving elements). We further estimated population receptive fields and used these to back-project the brain activity during stimulus perception into visual space via a searchlight procedure. Global perception resulted in non-topographic suppression of responses in lower visual cortex accompanied by wide-spread enhancement in higher object-sensitive cortex. However, follow-up experiments indicated that higher object-sensitive cortex is suppressed if global perception lacks shape grouping, and that grouping-related suppression can be diffusely confined to stimulated sites once stimulus size is reduced. These results speak against a rigid *between-area* response amplitude code acting as a generic grouping mechanism and point to a *within-area* response amplitude code mediating the perception of figure and ground.

## 1. Introduction

Perceptual grouping binds together local image elements into global and coherent objects and segregates them from other objects in our visual field including the background (Houtkamp, 2011; Roelfsema, 2006). This enables object recognition and tracking even if visual input is fragmented across space and time (Anderson & Sinha, 1997; Anstis & Kim, 2011; Lorenceau & Shiffrar, 1992), such as when we perceive a vehicle passing behind a row of trees. However, despite its ubiquity in everyday life, it remains unclear how perceptual grouping is represented in the visual brain.

A plethora of studies in monkeys suggests that information about figure-ground organization is represented in lower and mid-tier visual areas. In particular, neurons in V1 and V4 respond more strongly to tilted elements belonging to a global shape as opposed to the background (Lamme, 1995; Poort et al., 2016, 2012). Likewise, V1 and V4 responses to elements grouped into contours are enhanced, whereas those to ungrouped background elements are suppressed (Chen et al., 2014; Gilad et al., 2013). Taken together, these findings indicate that the monkey visual system draws upon a *within-area* response amplitude code to mediate figure-ground segregation.

Do similar mechanisms exist in humans? Although a series of (early) functional magnetic resonance imaging (fMRI) studies addressed this question (e.g., Altmann et al., 2003; Scholte et al., 2008; Seghier et al., 2000), their analyses techniques often lacked the spatial sensitivity to quantify retinotopically-constrained response amplitude codes. More recently, however, Kok & de Lange (2014) combined standard fMRI recordings and population receptive field (pRF) modeling (Dumoulin & Wandell, 2008) to investigate the topographic profile of V1 and V2 activity to illusory Kanizsa shapes in much greater detail. When compared to non-illusory control stimuli, activity to Kanizsa shapes increased, whereas activity to the illusion-inducing elements decreased, while background activity remained unchanged. Another topographic fMRI study reported ground-suppression in V1 (and also V2) without figure-enhancement for structure-from-asynchrony textures vs unstructured control stimuli (Likova & Tyler, 2008). Thus, here too, a within-area response amplitude mechanism emerges in lower visual areas, distinctively labelling multiple objects including the background.

The interpretation of these and similar studies is, however, complicated by the fact that changes in perception always went hand in hand with changes in the physical properties of the stimulus. This makes it impossible to determine unequivocally the source of such activity modulations. Bistable stimuli, for which our perception alternates between two mutually exclusive states without changes in the physical properties of the stimulus, provide a way to circumvent this issue. A very elegant bistable stimulus that allows for the investigation of perceptual grouping mechanisms in dynamic occluded scenes – where object tracking is often required – has been used by Fang et al. (2008) and Murray et al. (2002). In their studies, participants underwent fMRI while viewing a translating *diamond stimulus* whose corners were occluded by three bars of the same color as the background. This stimulus could either be perceived as four individual segments translating vertically out-of-phase and thus incoherently (*local, no-diamond* percept, Figure 1, A.) or as a diamond shape translating horizontally in-phase behind occluders and thus coherently (*global, diamond percept*; Figure 1, B., and Inline Supplementary Video 1). When participants experienced the global compared to the local percept, a striking pattern of results was observed: a reduction of activity in V1 (and also V2) accompanied by an increase of activity in the lateral-occipital complex (LOC) – a brain region known to respond more strongly to images of intact objects and shapes than a scrambled version thereof (e.g., Grill-Spector et al., 1998; Malach et al., 1995). Notably, this response pattern has recently been replicated (Grassi et al., 2018).

**Figure 1.**
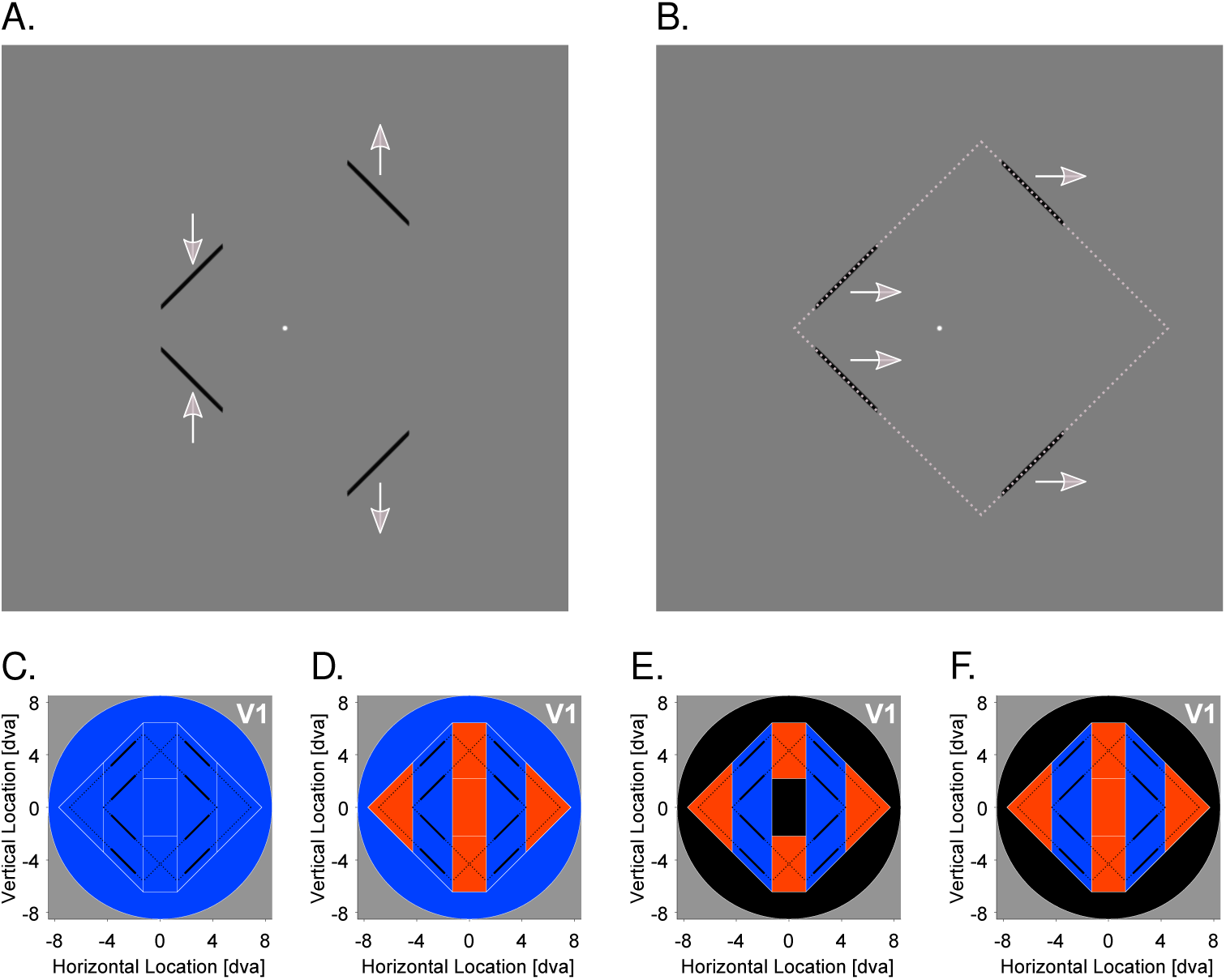
Diamond experiment. Example frames of the diamond stimulus and potential response amplitude profiles when the global percept is contrasted to the local one. **A.** Local, no-diamond percept. Here, the diamond stimulus was perceived as four individual segments oscillating vertically and incoherently with the segments on the left/right moving towards/away from one another, respectively, or vice versa (not shown). **B.** Global, diamond percept. Here, the four segments were grouped together and perceived as a diamond shape oscillating horizontally and coherently behind three occluders. The gray dashed frame denotes the inferred (but occluded) contours during the global state. The gray arrows indicate the perceived movement direction of the diamond stimulus. Only in the global state, the perceived and physical movement direction coincided. **C.** Previously suggested response amplitude profile. The whole visual field is suppressed. **D.** Hypothesized response amplitude profile. The segments and background region are suppressed whereas the corners and center regions are enhanced. **E.** Response amplitude profile when the segments, corners, and center region are predicted during the global state. The segments region is suppressed (due to a match between bottom-up input and higher-level feedback), the corners region enhanced (due to a mismatch between bottom-up input and higher-level feedback), and activity in the background and center region unchanged. **F.** The same as E., but if the whole diamond shape is predicted during the global state. The center region is now also enhanced. Black lines represent the extreme positions of the diamond stimulus. Black solid lines denote the visible ungrouped diamond segments (local, no-diamond percept). Black dashed lines additionally illustrate the inferred but invisible diamond shape when the segments were grouped together (global, diamond percept). White lines denote different visual field portions. Blue areas: Suppressive effects. Red areas: Enhancement effects. Black areas: No effect.

At first sight, such a *between-area* response amplitude mechanism reflects exactly the type of relationship proposed by *hierarchical predictive coding* models (e.g., Clark, 2013; Mumford, 1992; Murray et al., 2004; Rao & Ballard, 1999). These models assume that lower visual areas flag an error whenever the predictive feedback from higher visual areas conflicts with the bottom-up input they receive. The general idea here is that when higher visual areas (e.g., the LOC) arrive at a global and coherent interpretation of a visual stimulus (e.g., the diamond shape behind occluders), the predictability of the bottom-up input is increased and thus the error signal attenuated. When the global diamond percept is then contrasted to the local no-diamond percept, a differential reduction of activity emerges in lower visual areas (e.g., V1).

As such, these models predict that the reduction in V1 activity for the global percept should be restricted to the retinotopic representation of the visible diamond segments (Figure 1, E. and F.). This prediction, however, seems difficult to reconcile with the finding that the suppressive effects in V1 for the diamond vs no-diamond percept extend well beyond stimulated sites (i.e., the visible diamond segments) into the remaining background region (Figure 1, C.; De-Wit et al., 2012). It is also incompatible with evidence showing that variations of the diamond stimulus result in increased (instead of decreased) V1 activity for the diamond vs the no-diamond percept (Caclin et al., 2012).

These discrepant results may be due to the coarse analyses techniques employed previously, precluding a more fine-grained inspection of topographic signatures underlying the perception of the diamond stimulus. The possibility remains, for instance, that V1 activity corresponding to the region within the diamond frame (i.e., the center) and/or the invisible parts (i.e., the occluded corners) increases, whereas activity corresponding to the more peripheral background is suppressed during the diamond state (Figure 1, D.). De-Wit et al. (2012) considered much of these sub-areas as back-ground region, although the center and corners region could, arguably, be treated as figure and/or contour regions too. Although this hypothesis argues against hierarchical predictive coding models (e.g., Mumford, 1992; Murray et al., 2004; Rao & Ballard, 1999) because there should be no activity modulations in the peripheral background region (Figure 1, E. and F.), it is compatible with the more general idea of a within-area response amplitude mechanism labeling different parts of a visual scene distinctively (e.g., Gilad et al., 2013; Kok & de Lange, 2014; Lamme, 1995). Interestingly, such a response pattern has recently been observed for another dynamic bistable global-local stimulus (Grassi et al., 2017).

Here, we combined standard fMRI measurements and pRF modeling (similar to Kok & de Lange, 2014) to test for more fine-grained within-area and also between-area response amplitude mechanisms mediating global object perception. In a first experiment, we mapped the retinotopic organization of participants’ cortices and estimated the pRF of each voxel in visual cortex. In three further experiments, we recorded brain activity whilst participants viewed the diamond stimulus or a set of non-ambiguous stimuli with similar motion features but stable shape information to test for the generalizability of our findings. We then used each voxel’s pRF to back-project the voxel-wise brain activity measured during stimulus perception into visual space via a searchlight procedure. This allowed us to directly read-out retinotopically-specific response amplitude codes along a large portion of the visual hierarchy.

## 2. Retinotopic mapping experiment

### 2.1. Methods

#### 2.1.1. Participants

All participants (*N_total_* = 11) of the three global object perception experiments took part in the retinotopic mapping experiment. We refer to these participants as P1-P11. They all had normal or corrected-to-normal visual acuity and gave written informed consent to partake in our experiments (see 3.1.1, 4.1.1, and 5.1.1 Participants for more details). If participants took already part in the retinotopic mapping experiment in the scope of another study in our laboratory, we reused these data. All experimental procedures were approved by the University College London Research Ethics Committee.

#### 2.1.2. Apparatus

Functional and anatomical images were collected using a Siemens Avanto 1.5 Tesla magnetic resonance imaging (MRI) scanner. To prevent obstructed view, we used a customized version of the standard 32 channel coil, where the front visor was removed, reducing the number of channels to 30. For one participant (P2), however, the structural images were acquired with the standard 32 channel coil. Key presses were recorded via an MRI-button box for right-handers. Stimuli were projected onto a screen (resolution: 1920 *×* 1080 pixels; refresh rate: 60 Hz; background color: gray) at the back of the MRI scanner bore and viewed via a head-mounted mirror (viewing distance: approximately 67-68 cm; stimulus dimensions are based on the latter value; note that the variance in exact head/eye position is typically greater than this range). A list of software and toolboxes used in all experiments can be found in Supplementary Table S1.

#### 2.1.3. Stimuli

The retinotopic mapping stimulus consisted of a simultaneous wedge-and-ring aperture (Figure S1 and Inline Supplementary Video 2) centered within a screen-bounded rectangle in back-ground gray. The wedge aperture was a sector (polar angle: 12*^◦^*) of a disk (diameter: 17.03 dva), moving clockwise or counterclockwise in 60 discrete steps during 1 cycle (1 step/s). Consecutive wedges overlapped by 50%. The ring aperture consisted of an expanding or contracting annulus whose diameters varied in 36 logarithmic steps during 1 cycle (1 step/s). The diameter of the inner circle (minimum: 0.48 dva) was 56-58% of that of the outer circle (maximum: 40.38 dva, extending beyond the screen dimensions). The diameter of any current circle (outer or inner) was 10-11% larger/smaller compared to the previous one.

The wedge-and-ring aperture was superimposed onto circular images (diameter: 17.03 dva) depicting intact natural and colorful scenes/objects or a phase-scrambled version thereof (*N_total_* = 456). The images and the wedge-and-ring aperture were centered around a central black fixation dot (diameter: 0.13 dva) that was superimposed onto a central disk (diameter: 0.38 dva). Within the resulting annulus surrounding the fixation dot, the opacity level of the gray background increased radially inwards in 12 equal steps (step size: 0.02 dva) from fully transparent (*α* = 0 %) to fully opaque (*α* = 100 %).

To support fixation compliance, a black polar grid (line width: 0.02 dva) at low opacity (*α* = 10.2 %) centered around the fixation dot was superimposed onto the screen. The polar grid consisted of 10 circles whose diameters were evenly spaced between 0.38 and 27.35 dva, and 12 radial lines evenly spaced between polar angles of 0*^◦^* and 330*^◦^*. The radial lines extended from an eccentricity of 0.13 to 15.14 dva.

#### 2.1.4. Procedure

The retinotopic mapping experiment consisted of 3 runs. Excluding the initial dummy interval (10 s; fixation dot and polar grid only), each run comprised 4 blocks. At the beginning of each block, the wedge-and-ring aperture was presented (90 s; 1.5 cycles of wedge rotation; 2.5 cycles of ring expansion/contraction), followed by a fixation interval (30 s; fixation dot and polar grid only).

The order of wedge and ring movement in each run was clockwise and expanding (block 1), clockwise and contracting (block 2), counterclockwise and expanding (block 3), or counterclockwise and contracting (block 4). Within each block, the type of carrier image (intact or phase-scrambled) alternated every 15 s with the first carrier image always being phase-scrambled in odd-numbered blocks and intact in even-numbered blocks. The carrier images themselves were switched every 500 ms and displayed 1-2 times in pseudorandomized order during each run. To avoid confounds due to the spatial distribution of low-level features, the images were always rotated with the orientation of the wedge aperture.

Participants had to fixate the fixation dot continuously and press a key whenever the dot turned red. Every 200 ms, with a probability of 0.03, the fixation dot underwent a randomized change in color for 200 ms (from black to red, green, blue, cyan, magenta, yellow, white, or remaining black). To also ensure attention on the wedge- and-ring aperture, participants were required to press a key whenever a Tartan image appeared. Due to technical issues, for one participant (P3), the last 10 volumes (part of the final 30 s fixation interval) were not acquired in one run. To account for this, we also eliminated the last 10 volumes in the remaining two runs for this participant before submitting the functional data to our preprocessing procedure.

#### 2.1.5. MRI acquisition

Functional images were acquired with a T2*-weighted multiband 2D echo-planar imaging sequence (Breuer et al., 2005) from 36 transverse slices centered on the occipital cortex (repetition time, TR = 1 s, echo time, TE = 55 ms, voxel size = 2.3 mm isotropic, flip angle = 75*^◦^*, field of view, FoV = 224 mm *×* 224 mm, no gap, matrix size: 96 *×* 96, acceleration = 4). Slices were oriented to be approximately parallel to the calcarine sulcus while ensuring adequate coverage of the ventral occipital and inferior parietal cortex. Anatomical images were acquired with a T1-weighted magnetization-prepared rapid acquisition with gradient echo (MPRAGE) sequence (TR = 2.73 s, TE = 3.57 ms, voxel size = 1 mm isotropic, flip angle = 7*^◦^*, FoV = 256 mm *×* 224 mm, matrix size = 256 *×* 224, 176 sagittal slices).

#### 2.1.6. Preprocessing

After removing the first 10 volumes of each run to allow for T1-related signals to reach equilibrium, functional images were bias-corrected for intensity inhomogeneity, realigned, unwarped, and coregistered to the anatomical image. The anatomical image was used to construct a surface model, onto which the preprocessed functional data were projected. For each vertex in the surface mesh, we created an fMRI time series in each run by identifying the voxel in the functional images that fell half-way between the vertex coordinates in the gray-white matter and the pial surface. Finally, each time series was linearly detrended and *z*-standardized.

#### 2.1.7. Data analysis

##### PRF estimation

The preprocessed time series for each vertex were averaged across runs. To estimate the pRF for each vertex, we then implemented a forward-modeling approach restricted to the posterior third of the cortex. Each pRF was modeled as a 2D isotropic Gaussian with four free parameters: *x*, *y*, *σ*, and *β*, where *x* and *y* denote the pRF center position in Cartesian coordinates relative to fixation, *σ* the size of the pRF, and *β* the amplitude of the signal. The pRF center position and size were expressed in dva. The estimation procedure was identical to our previous studies (Moutsiana et al., 2016; van Dijk et al., 2016). The resulting parameter maps were modestly smoothed with a spherical Gaussian kernel (FWHM = 3 mm; for experiment-specific smoothing procedures of pRF and response data, see 3.1.7 Data analysis). Note that vertices with a very poor goodness-of-fit (*R*^2^ *≤* .01) were removed prior to smoothing.

##### Delineation of visual areas

Using the smoothed color-coded maps for eccentricity and polar angle projected onto the surface model of each hemisphere, we manually delineated V1-V3, V3A, V3B, LO-1, LO-2 (see all Wandell et al., 2007), V4, VO-1, and VO-2 (see all Winawer & Witthoft, 2015). Polar angle reversals served as primary indicator for identifying boundaries between visual areas (Engel et al., 1997; Sereno et al., 1995). Example maps used for back-projection purposes (see 3.1.7 Data analysis) including all delineations can be found in Supplementary Figure S1 (C. and D.).

For all data analyses, the quarterfield delineations of each hemisphere were merged and areas V3B, LO-1, LO-2, VO-1, and VO-2 combined into a larger complex we label the *ventral-lateral occipital complex* (VLOC). These sub-areas tended to show increased activation for intact vs phase-scrambled images (Supplementary Figure S1, E.), ensuring the functional validity of the VLOC as an object-sensitive complex. To this end, we performed a voxel-wise general linear model (GLM) for each participant on the preprocessed fMRI data from the retinotopic mapping experiment. The GLM comprised a constant boxcar regressor for each carrier type (intact vs phase-scrambled), convolved with a canonical hemodynamic response function. The fixation intervals were modeled implicitly and the obtained realignment estimates used as nuisance repressors. We applied Restricted Maximum Likelihood estimation with a first order autoregressive model, a high-pass filter (HPF) of 155 s, and implicit masking (threshold: 0.8). The voxel-wise differential beta values resulting from the GLM were then projected onto the surface model and smoothed moderately with a spherical Gaussian kernel (FWHM = 3 mm). Note that values flagged by implicit masking were discarded from smoothing and any subsequent visualizations. Similar functional localization procedures were applied previously to localize the LOC (e.g., De-Wit et al., 2012; Fang et al., 2008; Grill-Spector et al., 1998), which does typically not fully include the VO subareas and is not based on retinotopic principles. We thus refrained from labeling our complex ‘LOC’.

Importantly, compared to V1-V3, the subareas of the VLOC are smaller with fewer vertices and a sparser distribution of pRFs around the vertical meridian and the peripheral visual field (Amano et al., 2009; Larsson & Heeger, 2006). Combining these areas into the VLOC thus ensured a more complete coverage of the visual field in each participant, which was the basis for subsequent data analyses.

## 3. Diamond experiment

### 3.1. Methods

#### 3.1.1. Participants

Five healthy participants (P1-P5; 1 male; age range: 20-37 years; all right-handed), including the authors DSS and SS, took part in the diamond experiment.

#### 3.1.2. Apparatus

Apart from the apparatus of the retinotopic mapping experiment, we used an EyeLink 1000 MRI compatible eye tracker system to record eye movement data of participants’ left eye.

#### 3.1.3. Stimuli

The bistable diamond stimulus (similar to De-Wit et al., 2012; Fang et al., 2008) comprised a black rhombus-shaped frame (size: 7.92 *×* 7.92 dva; line width: 0.16) located around a white central fixation dot (diameter: 0.16 dva). Three vertical rectangles displayed in background color occluded the corners of the diamond stimulus. The middle rectangle (size: 3.75 *×* 17.03 dva) was centered around the fixation dot. The left and right rectangles (size: 22.84 *×* 17.03 dva, respectively) were centered vertically with their vertical line of symmetry coinciding with the left and right edges of the screen, so that the visible segments of the diamond had a length of 2.61 dva. When the diamond stimulus was centered around fixation, its corners were located at 5.6 dva eccentricity. The movement of the diamond followed a horizontal sine wave (*A* = 1.29 dva, *f* = 0.5 Hz, *ω* = 3.14, *φ* = 0).

The diamond display evoked two alternating and mutually exclusive perceptual states: a *local* percept of four individual segments translating vertically out-of-phase and thus incoherently (*no-diamond*; Figure 1, A.) or a *global* percept of an inferred diamond shape translating horizontally in-phase behind three occluders and thus coherently (*diamond*; see all Figure 1, B., and Supplementary Video 1).

#### 3.1.4. Procedure

The diamond experiment comprised 1 practice run (not analyzed) and 5 experimental runs. Experimental runs started with a background-only dummy interval (10 s). Next, an initial fixation interval (15 s) was presented, followed by the diamond display (400 s) and a final fixation interval (15 s). Except for the dummy interval, the fixation dot was continuously presented.

Participants were required to fixate the fixation dot continuously. During the diamond interval, they indicated their current percept via pressing a key assigned to their right index finger (diamond) or right middle finger (no-diamond). Except for the first percept in any given run, participants had to indicate perceptual switches only, but were allowed to press any key again if they lost track. During each run, participants’ eye position and pupil size were recorded at 60 Hz. Prior to scanning, all participants were tested behaviorally in a separate session outside the scanner to ensure they could clearly perceive both perceptual states and spent a roughly equal amount of time in either. Three recruited participants were unable to do so and hence replaced.

#### 3.1.5. MRI acquisition

Functional images were acquired with the same sequence as in the retinotopic mapping experiment.

#### 3.1.6. Preprocessing

The preprocessing was identical to the retinotopic mapping experiment using the same structural image.

#### 3.1.7. Data analysis

##### Searchlight back-projection

To explore intra- and also between-area response amplitude mechanisms, we first performed a voxel-wise GLM on the preprocessed data (HPF: 128 s). We used a variable epoch boxcar regressor (Grinband et al., 2008) for each perceptual state (diamond or no-diamond) as well as the period from the onset of the diamond display until participants’ first key press. The variable epochs for each perceptual state were the same as in the analysis of perceptual durations (see Supplementary material, 1.1.1 Data analysis). In all other respects (e.g. estimation procedure and nuisance regressors), the GLM was identical to the one specified for the retinotopic mapping experiment.

We computed the following contrasts of interest: *diamond vs fixation*, *no-diamond vs fixation*, and *diamond vs no-diamond*. The first two contrasts allowed us to verify the validity of our searchlight back-projection approach. Based on previous research on the positive and negative BOLD signal (Fracasso et al., 2018; Goense et al., 2012; Shmuel et al., 2002, 2006), we expected an increase of activity in the area within which the visible diamond segments moved and a decrease in non-stimulated sites, especially in lower visual areas (V1/V2), where pRF size is small (e.g., Alvarez et al., 2015; Amano et al., 2009; Dumoulin & Wandell, 2008; van Dijk et al., 2016). The contrast diamond vs no-diamond corresponded to analyses applied in prior studies involving the diamond stimulus (e.g., De-Wit et al., 2012; Fang et al., 2008). Based on the study by Fang et al. (2008) and De-Wit et al. (2012), we expected decreased activity in the area within which the diamond segments moved. However, we had no clear expectations as to how the remaining visual field would behave due to the coarser analyses techniques applied previously (De-Wit et al., 2012), evidence from figure-ground studies (Chen et al., 2014; Gilad et al., 2013; Gilad & Slovin, 2015; Kok & de Lange, 2014; Lamme, 1995; Poort et al., 2012, 2016), and findings showing increased activity for the diamond vs no-diamond percept (Caclin et al., 2012).

The voxel-wise differential beta values from the GLM were subsequently projected onto the surface model. Both the raw pRF data and the differential beta estimates were then modestly smoothed in an identical fashion using a spherical Gaussian kernel (FWHM = 3 mm). Vertices whose pRF estimates showed a very poor goodness-of-fit (*R*^2^ *≤* .01) or artifacts (*σ* or *β ≤* 0) were removed prior to smoothing. Vertices flagged by implicit masking were likewise discarded from smoothing as well as any subsequent analyses. We then used the delineations for each visual area and hemisphere from the retinotopic mapping experiment to extract pRF estimates and differential beta estimates of vertices falling within their spatial extent and pooled them across hemispheres for each participant. Vertices whose pRF estimates showed poor goodness-of-fit (*R*^2^ *≤* .05), and/or eccentricities outside the stimulated retinotopic mapping area (*≥* 8.5 dva) were discarded.

Subsequently, we defined a mesh grid (size: 17 *×* 17 dva) covering the stimulated retinotopic mapping area. The grid point coordinates were separated from one another by 0.1 dva in both the horizontal and vertical dimension (range: −8.5-8.5 dva, respectively). Next, a circular searchlight (radius: 1 dva) was passed through visual space by translating its center point from one grid point to the next. All vertices whose pRF center position fell into a given searchlight at a particular location were then identified. The differential beta estimates corresponding to the set of vertices within a given searchlight were summarized as a *t*-statistic by performing a one-sample *t*-test against 0. This way, we were able to account for the different numbers of vertices in each searchlight. *T-*statistics based on a single vertex/no vertices were set to 0. Importantly, *t*-statistics were only used as descriptive measure here. Of note, this searchlight procedure automatically normalizes the input data into a standard space as defined by the mesh grid.

For the vertices within a given searchlight, we derived the inverse Euclidean distance of their pRF center position from the respective searchlight center, normalized by the searchlight radius. These normalized vertex-wise weights were summed up searchlight-wise, resulting in summary weights where higher values reflect a higher number of vertices within a given searchlight as well as vertices with a pRF center position closer to the searchlight center. The summary weights were then normalized via dividing them by the 25*^th^* percentile of the resulting distribution of summary weights. Normalized summary weights *>* 1 were set to 1. Summary weights based on a single vertex were set to 0. Using the grid point coordinates, the resulting *t*-statistic maps were visualized as a heatmap. The color saturation of the heatmap was calibrated using the normalized summary weights, so that a higher saturation reflected a higher normalized summary weight.

The searchlight back-projections were obtained for each visual area and contrast of interest by pooling the data from all participants (after participant-wise smoothing). The pooling of data across participants improved the precision of searchlight back-projections because vertices from different participants complemented one another and covered the visual field more completely. Due to insufficient visual field coverage in V3A and V4 in each participant, we excluded these areas from the searchlight and all subsequent analyses.

##### Representational similarity analysis of searchlight back-projections

To explore the impact of each participant’s data set on the pooled searchlight back-projections, we performed a representational similarity analysis (Kriegeskorte, 2008). To this end, we first conducted a leave-one-subject-out (LOSO) analysis by repeating the search-light back-projections analysis whilst iteratively leaving out one participant. We then determined the dissimilarity (1-Spearman correlation) between the LOSO and the pooled back-projection matrices. Moreover, to assess the similarity structure more comprehensively, we also determined the dissimilarity between the individual (i.e., participant-wise) and the LOSO or pooled back-projections matrices. Importantly, for each back-projection pair, *t*-statistics based on a single vertex/no vertices were removed from both matrices prior to calculating the dissimilarity measure.

To visually summarize the dissimilarity structure, the resulting square matrices of dissimilarities (with zeros along the diagonal) were projected onto a 2D ordination space via Kruskall’s (1964a; 1964b) non-metric multidimensional scaling (NMDS) using monotone regression (criterion: stress). The final solution (based on 100 random starts) was centered, rotated via principal component analysis, and scaled to the range of the input dissimilarity measures. The lower the dissimilarity between two back-projection matrices, the closer they should be located in the 2D ordination space. Accordingly, if the pooled back-projections are representative of the whole study sample, the LOSO and individual back-projections should tightly cluster around or coincide with them.

### 3.2. Results

#### 3.2.1. Searchlight back-projections

Figure 2 depicts the searchlight back-projections for the pooled data per visual area and contrast of interest. When comparing the diamond or no-diamond percept to fixation, activity increased in the area within which the visible diamond segments moved. This pattern was fairly focal in V1 with suppressed differential activity in non-stimulated sites, but became more diffuse in V2, V3, and the VLOC.

**Figure 2.**
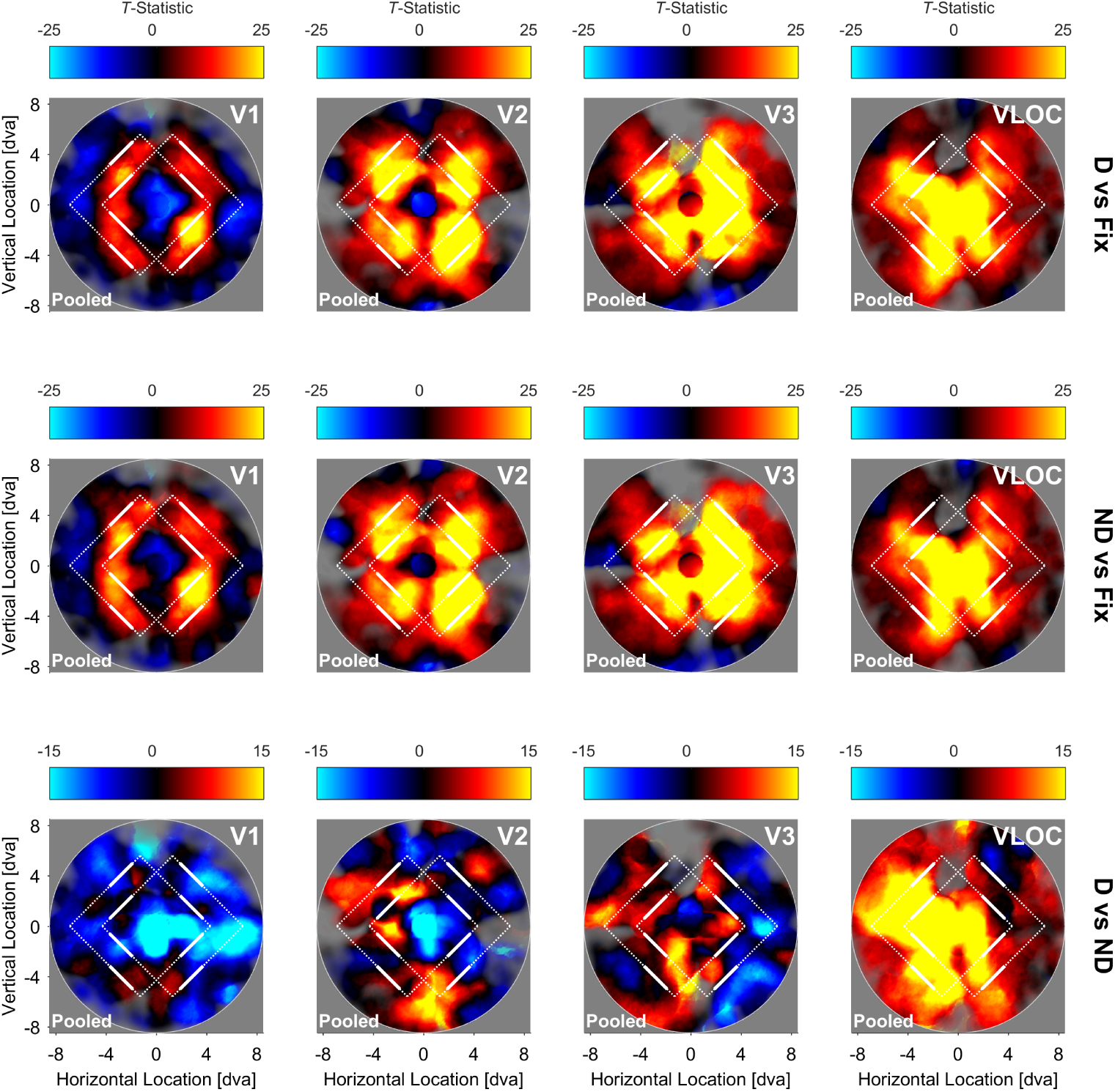
Diamond experiment. Searchlight back-projections of differential brain activity as a function of contrast of interest and visual area. *T-*statistics surpassing a value of ± 25 (first and second row) or ± 15 (third row) were set to that value. The saturation of colors reflects the number of vertices in a given searchlight plus their inverse distance from the searchlight center. White lines represent the extreme positions of the diamond stimulus. White solid lines denote the visible ungrouped diamond segments. White dashed lines additionally illustrate the inferred but invisible diamond shape when the segments were grouped together. D = Global, diamond percept. ND = Local, no-diamond percept. Fix = Fixation baseline. VLOC = Ventral-and-lateral occipital complex. Pooled = Data pooled across all 5 participants.

For the contrast diamond vs no-diamond, we observed a wide-spread suppression of activity in V1, particularly along the horizontal meridian. Although V2 and V3 showed similar suppressive effects, these were less extensive and intermixed with distinct opposite effects. There was also no clear indication of a suppression streak along the horizontal meridian. Finally, unlike V1-V3, the contrast diamond vs no-diamond showed a wide-spread increase of activity in the VLOC.

#### 3.2.2. Representational similarity of searchlight back-projections

Figure 3 depicts the NMDS solution for dissimilarities calculated between the individual, pooled, and LOSO searchlight back-projections, separately for each contrast of interest and visual area. The corresponding representational dissimilarity matrices can be found in Supplementary Figure S3.

**Figure 3.**
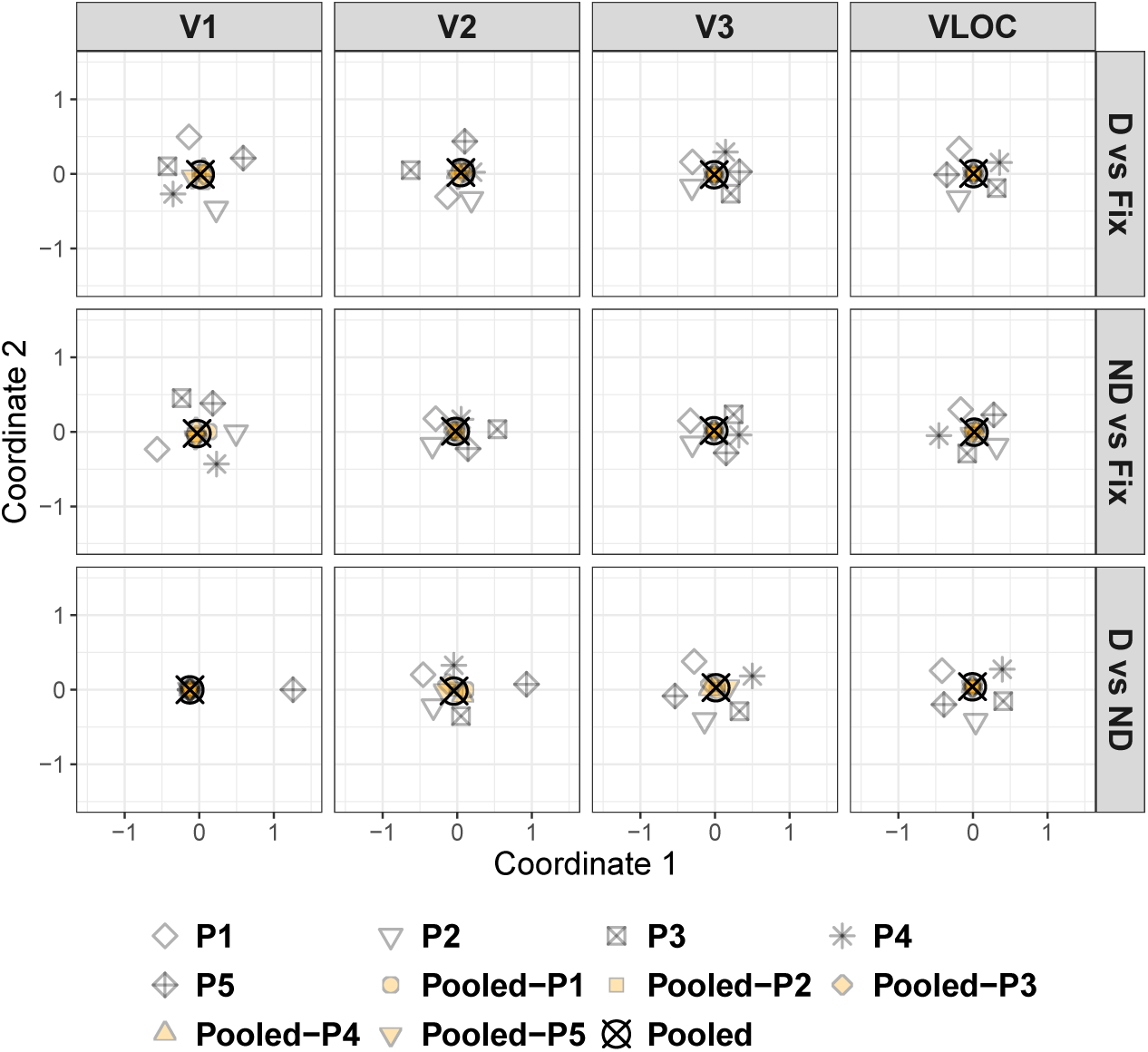
Diamond experiment. Non-metric multi-dimensional scaling of the dissimilarities from Figure S3 as a function of contrast of interest and visual area. D = Global, diamond percept. ND = Local, no-diamond percept. Fix = Fixation baseline. VLOC = Ventral-and-lateral occipital complex. P1-P5 = Participant 1-5. Pooled = Data pooled across all 5 participants. Pooled-P1-Pooled-P5 = Data pooled across 4 participants with 1 participant left out (as indicated by the suffix). LOSO = Leave-one-subject-out.

For all visual areas and contrasts, the LOSO back-projections essentially coincided with the pooled back-projections, highlighting a low degree of dissimilarity. Thus, the pooled back-projections do not seem to be driven by a single participant. The individual back-projections clustered around the pooled ones in a circular fashion, but less tightly than the LOSO back-projections, suggesting a higher degree of dissimilarity. Strikingly, for the contrast diamond vs no-diamond in V1 and V2, the back-projection pattern for P5 was located far away from the remaining ones, indicating a high degree of dissimilarity (see all Figure 3). Indeed, when examining the representational dissimilarity matrices directly (Figure S3), it becomes evident that the back-projections for P5 in V1 and V2 show a pattern opposite to the other participants.

### 3.3. Discussion

Here, we explored within- and between-area response amplitude codes in human visual cortex underlying global object perception. Participants viewed a bistable diamond stimulus that was either perceived as four individual segments moving vertically and incoherently (local, no-diamond percept) or a diamond shape drifting horizontally and coherently behind occluders (global, diamond percept).

When contrasting either the diamond or no-diamond percept to fixation, our searchlight back-projections revealed enhanced activity in cortical sites stimulated by the visible diamond segments. This differential increase was concise in V1 along with reduced activity in non-stimulated sites, but became more widespread in V2, V3, and the VLOC. We therefore replicate previous work on stimulus-evoked retino-topic activation and background suppression in visual cortex (Fracasso et al., 2018; Goense et al., 2012; Shmuel et al., 2002, 2006). Our findings furthermore comply with predictions based on between-area differences in pRF size (Alvarez et al., 2015; Amano et al., 2009; Dumoulin & Wandell, 2008; van Dijk et al., 2016). Specifically, given that pRF size is larger in higher visual areas, there is a greater number of peripherally located pRFs encoding the visible diamond segments, resulting in a more diffuse topographic representation. In sum, these results confirm our expectations and validate our searchlight back-projection approach.

When we directly compared the diamond to the no-diamond percept, our search-light analysis indicated a large-scale suppression of activity in V1 along with tendentially less extensive suppressive effects in V2 and V3. This global dampening effect speaks against the idea of a within-area response amplitude mechanism labelling different portions of the diamond display distinctively to mediate global object perception (Chen et al., 2014; Gilad et al., 2013; Gilad & Slovin, 2015; Grassi et al., 2017; Kok & de Lange, 2014; Lamme, 1995; Likova & Tyler, 2008; Poort et al., 2012, 2016). Critically, however, it echoes prior reports of retinotopically-unspecific deactivation during the diamond vs no-diamond percept and an attenuation of these effects in V2/V3 (De-Wit et al., 2012).

In contrast, there was a wide-spread enhancement of activity in the VLOC for the diamond compared to the no-diamond percept. This mirrors previous studies on the diamond stimulus identifying the LOC as a source for modulatory feedback in lower visual areas (Fang et al., 2008; Murray et al., 2002). This idea is corroborated by a large body of work highlighting the sensitivity of LOC responses to global shape and intact objects even under occlusion conditions (Grill-Spector et al., 1999; Hegdé et al., 2008; Lerner et al., 2002, 2004; Malach et al., 1995; Vinberg & Grill-Spector, 2008). Moreover, given that visual stimulation was identical in the diamond and no-diamond percept, the universal deactivation we observed in lower visual cortex cannot be attributed to physical stimulus differences (Dumoulin & Hess, 2006) and was thus likely subject to top-down modulation.

However, it is unclear whether the inverse relationship between the VLOC/LOC and lower visual cortex we and others quantified (Fang et al., 2008; Grassi et al., 2018; Murray et al., 2002) can be regarded as a generic perceptual grouping mechanism operating irrespective of shape perception. Recent evidence suggests, for instance, that activity in the LOC also decreases for intact vs scattered objects with abolished inter-part relations (Margalit et al., 2017) as it is the case during the no-diamond percept. In order to address this question, our third experiment used a non-ambiguous stimulus consisting of four circular apertures, each carrying a random dot kinematogram (RDK). In the local condition, the RDKs translated vertically and incoherently. In the global condition, however, they moved horizontally and coherently and could thus be grouped together without forming a hybrid shape. These conditions closely echoed the motion features of the diamond stimulus whilst keeping shape information (i.e., the four circular apertures) constant and allowing for perceptual grouping. If the between-area response amplitude mechanism between the VLOC/LOC and lower visual cortex indeed constitutes a generic grouping mechanism, we should be able to conceptually replicate the findings from our diamond experiment.

## 4. Dots experiment

### 4.1. Methods

#### 4.1.1. Participants

The authors DSS and SS as well as 3 other healthy participants (P1, P2 and P6-P8; 1 male; age range: 24-38 years; 1 left-handed) partook in this experiment.

#### 4.1.2. Apparatus

All apparatus were identical to the diamond experiment although the viewing distance to the head-mounted mirror was approximately 67 cm here as this facilitated the use of the eye tracker.

#### 4.1.3. Stimuli

The dots stimulus comprised four circular apertures through which a random dot kinematogram (RDK), that is, a field (size: 2.85 *×* 2.85 dva) of moving black dots (diameter: 0.11 dva) was presented. The apertures were generated by removing all dots falling outside or on the edge of a circle (diameter: 2.85 dva) centered within the dots field. The aperture centers were positioned at the corners of a square (size: 5.69 *×* 5.69 dva) centered around a white central fixation dot (diameter: 0.16 dva). The dots of each aperture had a density of 12.33 dots/dva^2^. All dots had a lifetime of 9 frames and were repositioned randomly within their field once they died. If the dots moved beyond the edge of their field, they were moved back by 1 field width. The position of a given dot at the beginning of each block was determined randomly as was the time a dot had already lived.

In the *global horizontal* condition, the dots in all apertures moved synchronously according to a horizontal sine wave (*A* = 1.31 dva, *f* = 0.5 Hz, *ω* = 3.14, *φ* = 0; Figure 4, B.). In the *local vertical* condition, they followed an identical but vertical sine wave with the dots in the bottom-right and top-left apertures moving anti-synchronously (*φ*_1_ = 0) relative to the dots in the top-right and bottom-left apertures (*φ*_2_ = *π*; Figure 4, A., and Inline Supplementary Video 3). The horizontal condition mimicked the perceived movement during the global diamond percept and enabled participants to group the 4 apertures together through the Gestalt principle of common fate similar to the diamond stimulus. The vertical condition mirrored the perceived movement during the local no-diamond percept. Notably, the number of apertures and shape information remained the same in both conditions.

**Figure 4.**
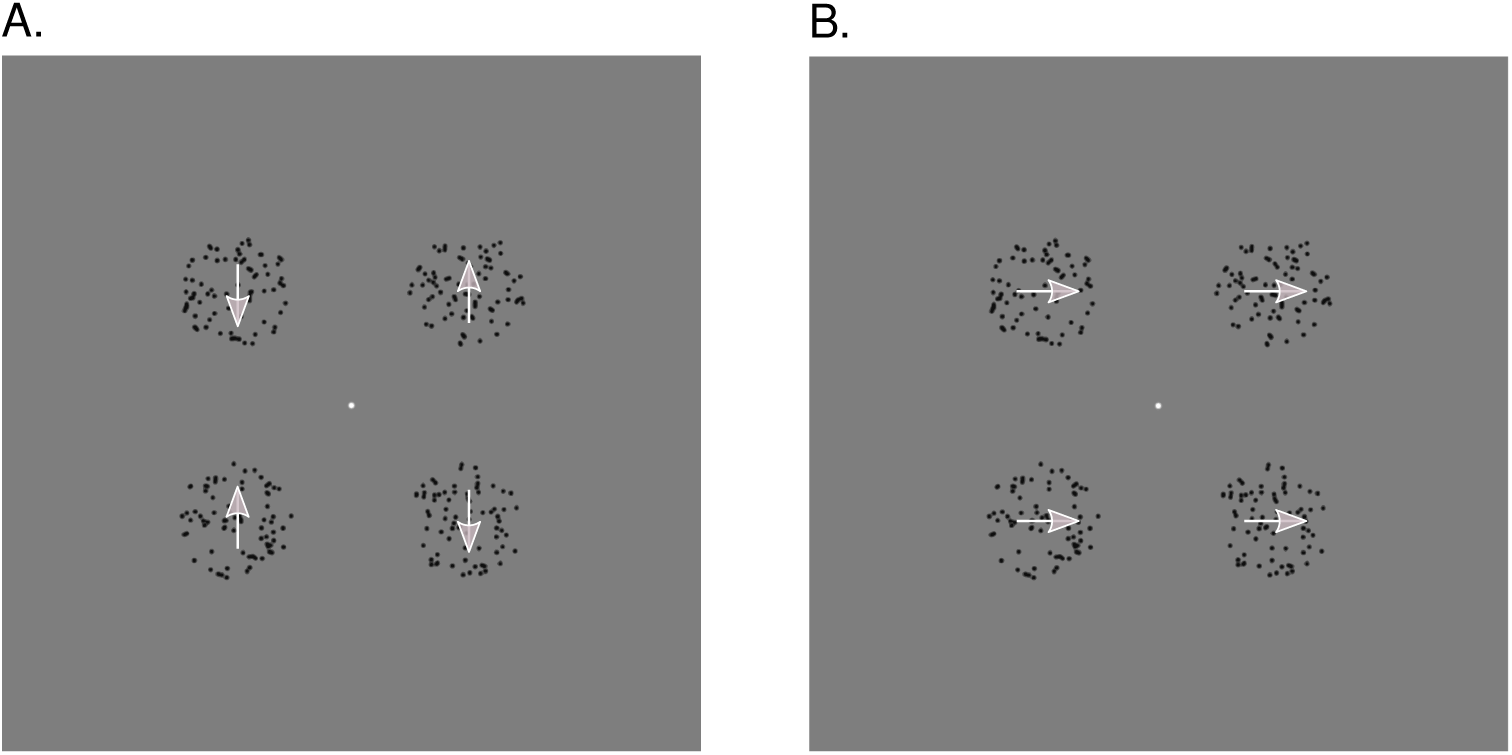
Dots experiment. Example frames of the dots stimulus. **A.** Local, vertical condition. Here, the dots oscillated vertically and incoherently with the dots in the left/right apertures moving towards/away from one another, respectively, or vice versa (not shown), so that the apertures were perceived as 4 individual elements. **B.** Global, horizontal condition. Here, the dots in all apertures oscillated horizontally and coherently, so that the apertures could be grouped together into a global Gestalt without forming a hybrid shape. Since this stimulus was non-ambiguous, the gray arrows naturally indicate the perceived and physical movement direction of the dots within the aperture.

#### 4.1.4. Procedure

The dots experiment comprised 8 experimental runs. Excluding the initial dummy interval (10 s without fixation dot), each run was split into 8 blocks. Within each block, a fixation interval (15 s) was presented followed by the dots stimulus (30 s) in either the vertical or horizontal condition. Within each run, the horizontal and vertical conditions were presented in an alternating fashion, starting with the vertical condition in uneven-numbered and the horizontal condition in even-numbered runs. At the end of each run, a final fixation interval (15 s) was displayed.

Participants were required to fixate the fixation dot continuously. In the dots interval, they indicated whenever the dots in one of the circular apertures flickered shortly (by changing their color to background gray for 200 ms) via pressing a key with their right index finger (left apertures) or right middle finger (right apertures). The number of flicker events per block was determined randomly but was always 3, 6, or 9 with a gap of at least 200 ms between consecutive flicker events. The aperture within which the flicker events occurred was determined randomly. Participants’ eye position and pupil size were recorded at 60 Hz.

#### 4.1.5. MRI acquisition

The MRI acquisition was as in the retinotopic mapping and diamond experiment.

#### 4.1.6. Preprocessing

The preprocessing was identical to the retinotopic mapping and diamond experiment. It is of note, however, that P7 moved more than other participants during the dots experiment. Moreover, for this participant, coregistration in the retinotopic experiment was also less ideal than for others. It is thus important to perform any analyses with and without this participant.

#### 4.1.7. Data analysis

##### Searchlight back-projection

The searchlight back-projection analysis was conducted in the same manner as in the diamond experiment with exceptions as follows. The voxel-wise GLM on the preprocessed data (HPF: 185 s) involved a constant epoch boxcar regressor for each condition (horizontal or vertical) and an event-related regressor for the onset of the flicker events. We calculated the following contrasts of interest: *horizontal vs fixation*, *vertical vs fixation*, and *horizontal vs vertical*. The contrasts horizontal or vertical vs fixation were equivalent to the contrasts diamond or no-diamond vs fixation, respectively. The contrast horizontal vs vertical mirrored the contrast diamond vs no-diamond.

##### Representational similarity of searchlight back-projections

The representational similarity analysis was conducted as in the diamond experiment.

### 4.2. Results

#### 4.2.1. Searchlight back-projections

Figure 5 shows the back-projected searchlight-based profiles pooled across participants for each visual area and contrast of interest. When comparing the horizontal or vertical condition to fixation, there was enhanced activity in areas carrying the RDKs. This pattern was spatially relatively precise in V1 with suppressive effects in the central and peripheral visual field, and became more wide-spread in V2, V3, and the VLOC.

**Figure 5.**
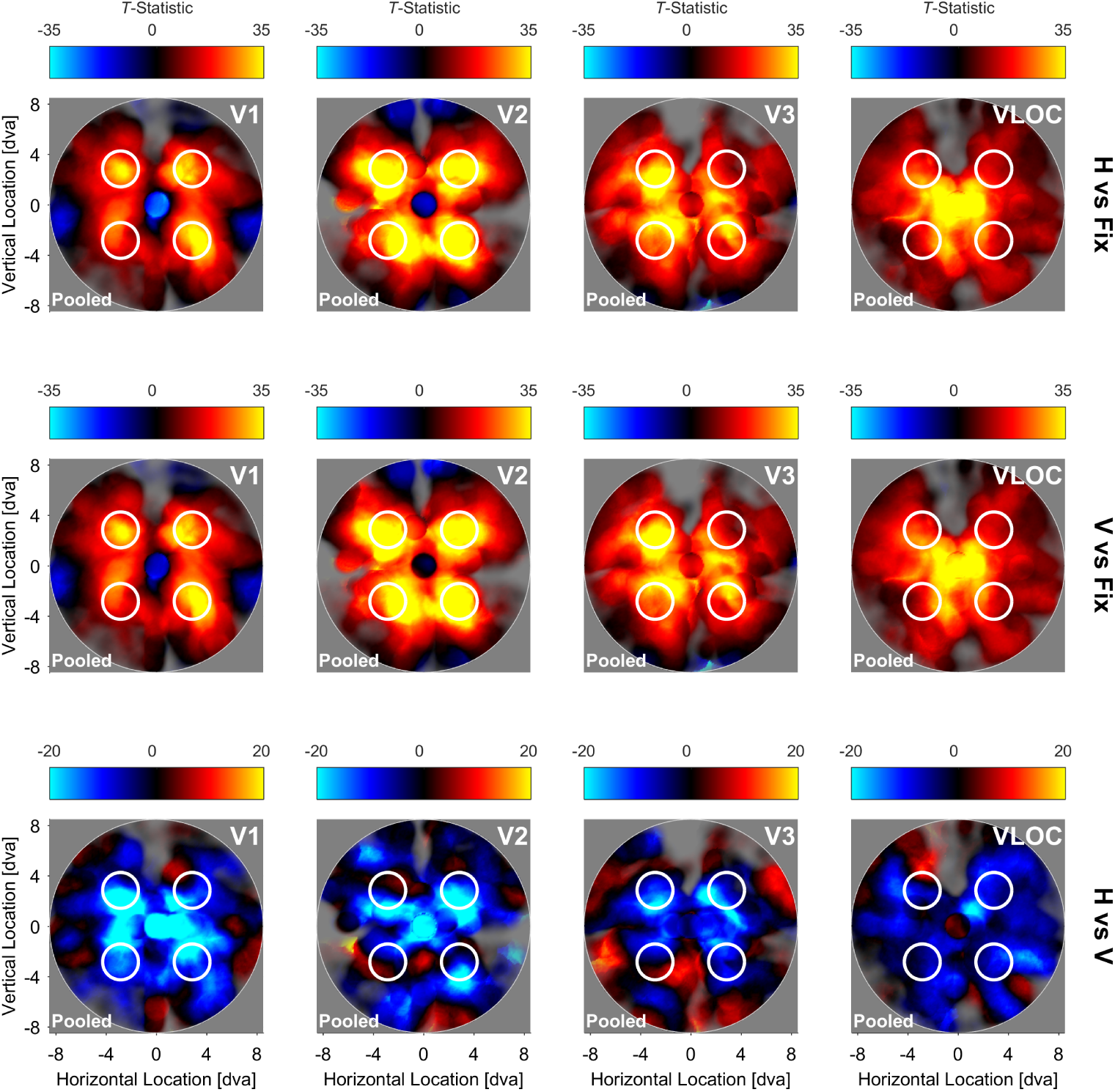
Dots experiment. Searchlight back-projections of differential brain activity as a function of contrast of interest and visual area. *T-*statistics surpassing a value of ± 35 (first and second row) or ± 20 (third row) were set to that value. The saturation of colors reflects the number of vertices in a given searchlight plus their inverse distance from the searchlight center. White lines represent the spatial extent of the circular apertures carrying the RDK. H = Global, horizontal condition. V = Local, vertical condition. Fix = Fixation baseline. VLOC = Ventral-and-lateral occipital complex. Pooled = Data pooled across all 5 participants. RDK = Random dot kinematogram.

For the direct comparison between the horizontal and vertical condition, we observed a fairly wide-spread deactivation across the whole visual field in all visual areas, occasionally intermixed with fairly focal opposite effects. These diffuse suppressive effects were particularly eminent around the central visual field and stimulated areas but not in the background area.

#### 4.2.2. Representational similarity of searchlight back-projections

Figure 6 illustrates the NMDS solution for the dissimilarities between the individual, pooled, and LOSO searchlight back-projections per contrast of interest and visual area. Supplementary Figure S4 shows the corresponding representational dissimilarity matrices.

**Figure 6.**
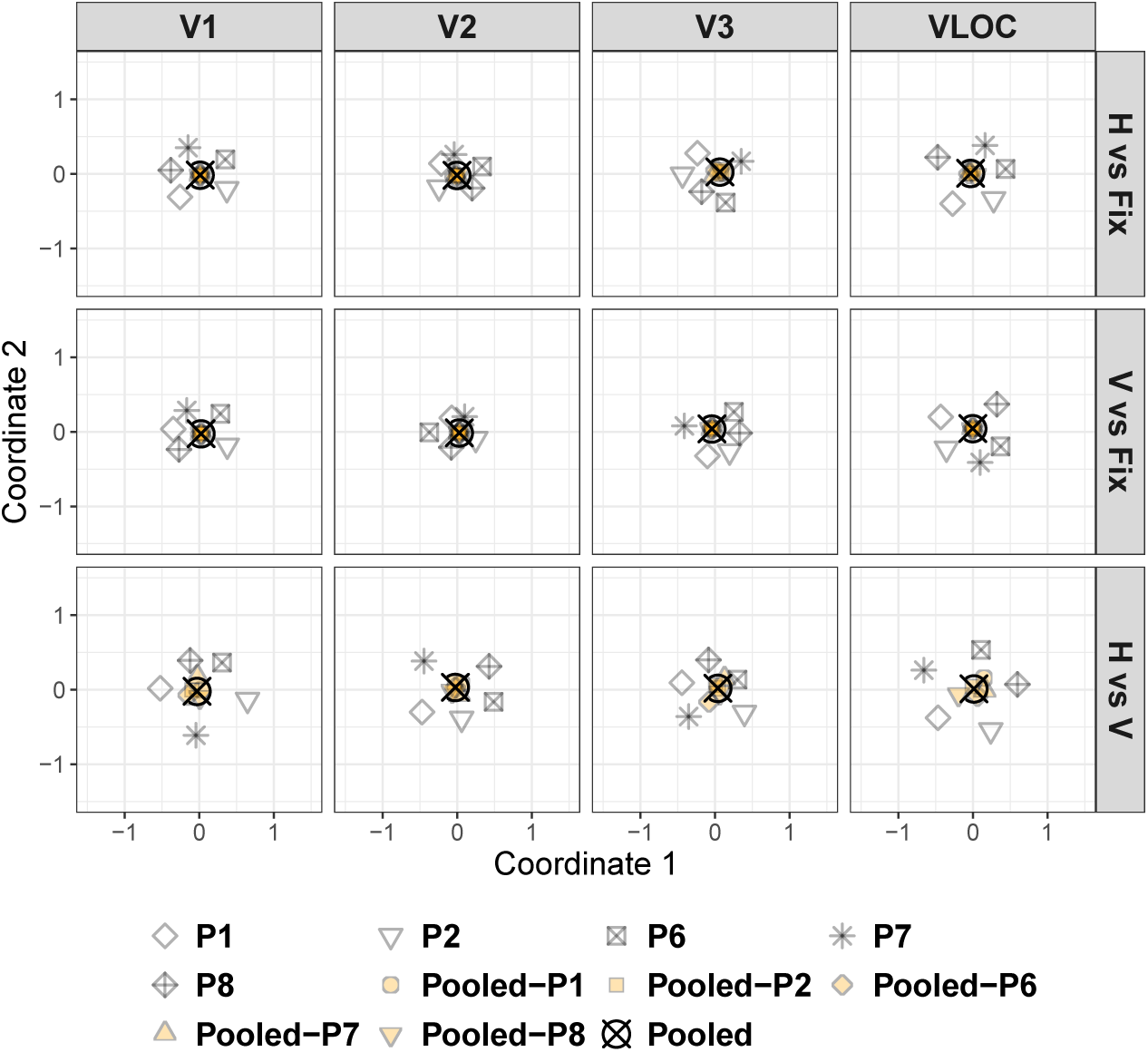
Dots experiment. Non-metric multi-dimensional scaling of the dissimilarities from Figure S4 as a function of contrast of interest and visual area. H = Global, horizontal condition. V = Local, vertical condition. Fix = Fixation baseline. VLOC = Ventral- and-lateral occipital complex. P1-P2 and P6-P8 = Participant 1-2 and 6-8. Pooled = Data pooled across all 5 participants. Pooled-P1-Pooled-P2 and Pooled-P6-Pooled-P8 = Data pooled across 4 participants with 1 participant left out (as indicated by the suffix). LOSO = Leave-one-subject-out.

The LOSO back-projections generally accorded well with the pooled ones, high-lighting a low degree of dissimilarity. As such, the pooled back-projections do not seem to be driven by single participants including P7 who moved more than other participants and for whom coregistration was difficult. The individual back-projections clustered circularly around the pooled ones, albeit less closely than the LOSO back-projections, indicating a higher degree of dissimilarity. This was particularly eminent for the contrast horizontal vs vertical in V1 and the VLOC (see all Figure 6). As the representational dissimilarity matrices indicate (Supplementary Figure S4), this pattern highlights the highly idiosyncratic nature of the individual back-projections.

### 4.3. Discussion

Here, we investigated between- and within-area response amplitude mechanisms related to the perception of a global Gestalt in an attempt to generalize the findings of our diamond experiment beyond shape perception. Participants viewed four apertures carrying random dots that moved either vertically and incoherently (local, vertical condition) or horizontally and coherently, allowing perceptual grouping into a global configuration (global, horizontal condition). These conditions echoed the global-local aspects of the diamond stimulus without varying in shape information. We hypothesized that if the between-area response amplitude mechanism between lower visual cortex and VLOC/LOC we and others observed (Fang et al., 2008; Grassi et al., 2018; Murray et al., 2002) indeed mediates global object perception per se, we should be able to conceptually replicate this relationship.

To validate our analysis procedures, we compared the horizontal or vertical condition to fixation. Our searchlight back-projections highlighted increased differential activity in physically stimulated sites and suppressive effects in non-stimulated sites. The spatial precision of this pattern was relatively high in V1 and decreased from V2 over V3 to the VLOC. Collectively, these results are in line with our diamond experiment and confirm the spatial sensitivity of our back-projection approach.

To generalize the findings of our diamond experiment, we compared the horizontal and vertical condition directly, revealing a diffuse pattern of suppressed differential activity across large portions of the visual field in all visual areas. The wide-spread deactivation in lower visual cortex is consistent with our previous diamond results. The diffuse deactivation in the VLOC, however, contradicts the idea that its previously established inverse relationship to lower visual cortex represents a between-area response amplitude mechanism mediating global object perception beyond shape perception.

An interesting additional finding is that V1 and V2 activity in the more peripheral back-ground area did not seem to be strongly suppressed for the horizontal relative to the vertical condition, but showed a tendency to remain unchanged or slightly enhanced. This could suggest that the dampening effects we observed are diffusely related to the stimulus and level out further in the periphery. Alternatively, this may be related to a comparably sparser distribution of pRFs in the background area along with a fairly large size and central presentation of the dots stimulus and thus relative undersampling of the background area. Consequently, the question arises as to whether the large-scale deactivation in lower visual cortex also occurs if the dots stimulus is smaller, e.g., confined to one visual field quadrant only. Critically, if this were not the case and the deactivation quadrant-specific and not present in the remaining visual field, this could be regarded as a diffuse instantiation of a within-area response amplitude mechanism. In our fourth experiment, we therefore essentially repeated the dots experiment, but moved the dots stimulus to the top-right visual field quadrant.

## 5. Dots quadrant experiment

### 5.1. Methods

#### 5.1.1. Participants

The author SS and 4 other healthy participants (P1, P6, and P9-P11; 1 male; age range: 20-36 years; all right-handed) participated in this experiment.

#### 5.1.2. Apparatus

All apparatus were identical to the dots experiment.

#### 5.1.3. Stimuli

The dots quadrant stimulus was identical to the dots stimulus except that the stimulus configuration was smaller and repositioned. Specifically, the dots field subtended 0.58 *×* 0.58 dva and the diameter of the circular apertures was thus 0.58 dva. The aperture midpoints were centered around the corners of a square with a size of 2.27 *×* 2.27 dva. The dots configuration was always presented in the top-right visual field quadrant. Its midpoint was located at a distance of 3.41 dva in the x- and y-direction from the center of the screen. The density of the dots in each aperture was 60.31/dva^2^ and thus higher than in the dots experiment. This way, we ensured that the movement of the dots was still clearly perceivable. As in the dots experiment, there was a local *vertical* (Figure 7, A.) and global *horizontal* condition (Figure 7, B., and Inline Supplementary Video 4).

**Figure 7.**
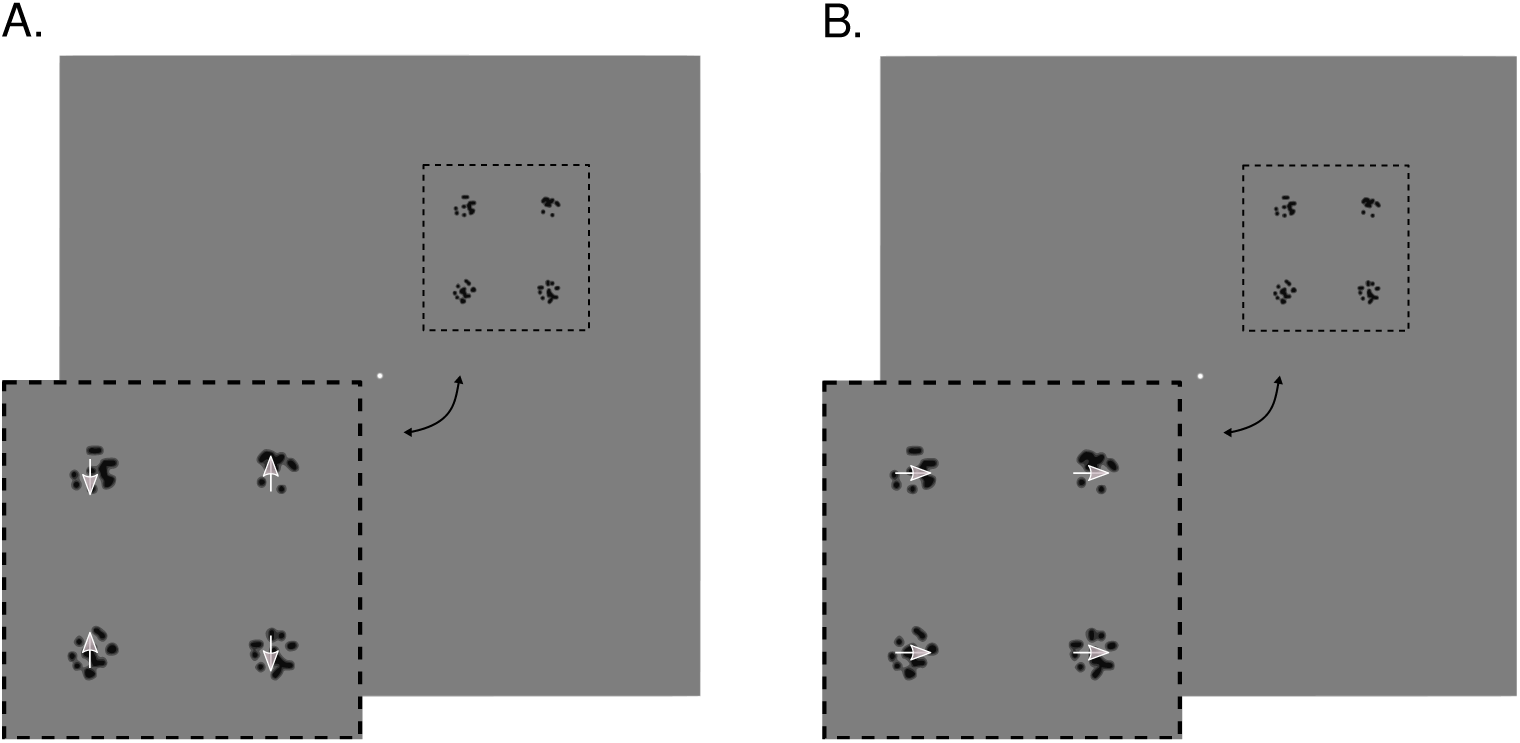
Dots quadrant experiment. Example frames of the dots quadrant stimulus. **A.** Local, vertical condition. Here, the dots oscillated vertically and incoherently with the dots in the leftmost/rightmost apertures moving towards/away from one another, respectively, or vice versa (not shown), so that the apertures were perceived as 4 individual elements. **B.** Global, horizontal condition. Here, the dots in all apertures oscillated horizontally and coherently, so that the apertures could be grouped together into a global Gestalt without forming a hybrid shape. Since this stimulus was non-ambiguous, the gray arrows naturally indicate the perceived and physical movement direction of the dots within the aperture. The dots quadrant stimulus was only presented in the top-right visual field quadrant. For reasons of visibility, we cut out the stimulus region to provide a zoomed-in view, as indicated by the black dashed lines and the black double-headed arrows.

#### 5.1.4. Procedure

The procedure of the dots quadrant experiment was the same as for the dots experiment, although here, participants were required to press their right index/middle finger when the dots of any of the leftmost/rightmost apertures flickered.

#### 5.1.5. MRI acquisition

The MRI acquisition was identical to the other experiments except that we additionally collected a rapid MPRAGE (TR ^1^= 1.150 s, TE = 3.6 ms, voxel size = 2 mm isotropic, flip angle = 7*^◦^*, FoV = 256 mm *×* 208 mm, matrix size = 128 *×* 104, 80 sagittal slices) to aid coregistration of the functional to the structural images if the structural image was acquired in a separate session.

#### 5.1.6. Preprocessing

The preprocessing was identical to all other experiments. However, if rerunning automated coregistration after manual registration failed, we performed a 2-pass-procedure where the functional images were first coregistered to the short MPRAGE and then to the long MPRAGE. Where necessary, this 2-pass-procedure was also applied to the retinotopic mapping data of a given participant.

#### 5.1.7. Data analysis

##### Searchlight back-projections and representational similarity of searchlight back-projections

The searchlight back-projection and representational similarity analysis were conducted in the same manner as in the dots experiment.

### 5.2. Results

#### 5.2.1. Searchlight back-projections

Figure 8 depicts the searchlight back-projection profiles for the pooled data as a function of visual area and contrast of interest. When contrasting the horizontal or vertical condition to fixation, our back-projection profiles highlighted enhanced activity in stimulated visual field portions. This differential enhancement was confined to the top-right visual field quadrant in V1 and V2 with suppressive effects in the remaining quadrants, but increasingly extended into the top-left and bottom-right quadrants from V3 to the VLOC.

**Figure 8.**
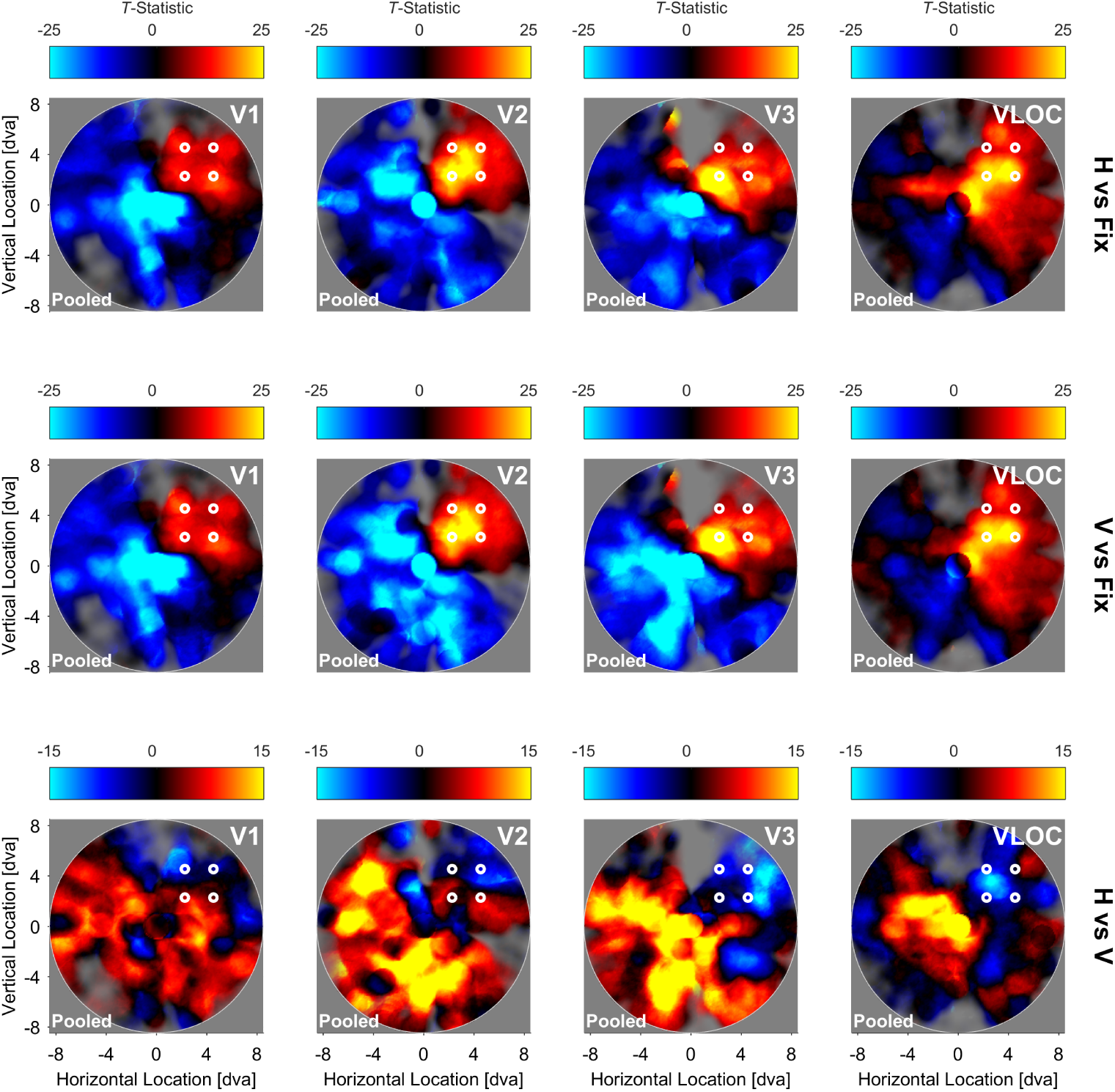
Dots quadrant experiment. Searchlight back-projections of differential brain activity as a function of contrast of interest and visual area. *T-*statistics surpassing a value of ± 25 (first and second row) or ± 15 (third row) were set to that value. The saturation of colors reflects the number of vertices in a given searchlight plus their inverse distance from the searchlight center. White lines represent the spatial extent of the circular apertures carrying the RDK. H = Global, horizontal condition. V = Local, vertical condition. Fix = Fixation baseline. VLOC = Ventral-and-lateral occipital complex. Pooled = Data pooled across all 5 participants. RDK = Random dot kinematogram.

For the contrast horizontal vs vertical, we observed a tendency for suppressive effects in stimulated areas of V1 and V2 and enhanced effects in the remaining visual field. In V3 and the VLOC, this pattern was much more pronounced and wide-spread.

#### 5.2.2. Representational similarity of searchlight back-projections

Figure 9 shows the NMDS solution for the dissimilarities calculated between the individual, pooled, and LOSO searchlight back-projections by contrast of interest and visual area. The corresponding representational dissimilarity matrices can be found in Supplementary Figure S5.

**Figure 9.**
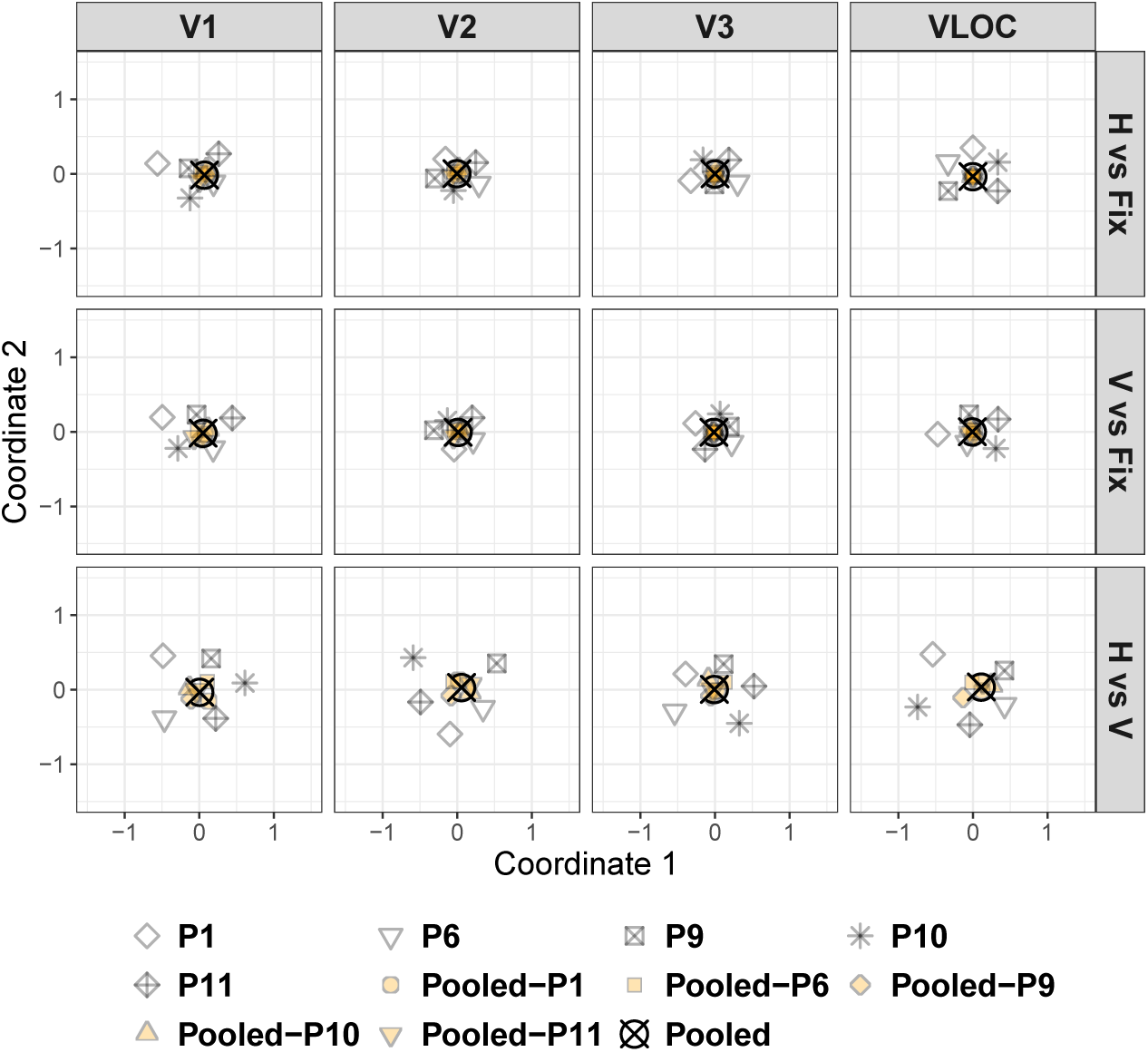
Dots quadrant experiment. Non-metric multi-dimensional scaling of the dissimilarities from Figure S5 as a function of contrast of interest and visual area. H = Global, horizontal condition. V = Local, vertical condition. Fix = Fixation baseline. VLOC = Ventral-and-lateral occipital complex. P1, P6, P9-P11 = Participant 1, 6, and 9-11. Pooled = Data pooled across all 5 participants. Pooled-P1, Pooled-P6, Pooled-P9-Pooled-P11 = Data pooled across 4 participants with 1 participant left out (as indicated by the suffix). LOSO = Leave-one-subject-out.

In virtually all cases, the LOSO back-projections coincided well with the pooled ones, suggesting a low degree of dissimilarity and thus speaking against an overly strong influence of single participants. The individual back-projections tended to cluster circularly around the pooled ones, albeit less tightly than the LOSO back-projections, highlighting a higher degree of dissimilarity. However, some individual back-projections were located far apart from one another or the pooled back-projections. This was particularly true for the contrast horizontal vs vertical in V1 and the VLOC (see all Figure 9). As confirmed by the representational dissimilarity matrices (Supplementary Figure S5), this structure is indicative of a fairly high degree of dissimilarity and with that inter-individual variability.

### 5.3. Discussion

Here, we tested for a diffuse instantiation of a within-area response amplitude mechanism related to parafoveal Gestalt perception. Participants viewed apertures filled with random dots in the top-right visual field quadrant. The dots moved either vertically and incoherently (local, vertical condition) or horizontally and coherently (global, horizontal condition). Based on the results of our dots experiment, we hypothesized that any suppression of activity might be diffusely related to the physical stimulus and thus the top-right visual field quadrant or bordering areas.

In line with our hypothesis, when contrasting the horizontal to the vertical condition, our searchlight back-projections revealed a trend for a reduction of activity near the stimulus location in V1 and V2 – a pattern that became more pronounced and wide-spread in V3 and the VLOC. Moreover, we observed an increase of activity in the remaining visual field in all visual areas. We therefore found evidence for a within-area enhancement-suppression mechanism mediating the perception of figure and ground, as previously established in macaques (Chen et al., 2014; Gilad et al., 2013; Gilad & Slovin, 2015; Lamme, 1995; Poort et al., 2012, 2016) and humans (Grassi et al., 2017; Kok & de Lange, 2014; Likova & Tyler, 2008).

The absence of clear suppressive effects in V1 and V2 (as compared to V3 and the VLOC) might be related to the functional architecture of the visual cortex, noisy voxels, and the size of the dots quadrant stimulus. Specifically, in lower visual areas, pRFs are smaller and with that the number of pRFs encoding the physical stimulus tendentially reduced (although not necessarily), resulting in diminished response gain. Consequently, noisy voxels are likely to have a more pronounced impact on searchlight-wise response amplitude quantifications. Moreover, stimulus-driven activity modulations tend to be weaker for smaller and more eccentric stimuli (Nasr et al., 2015) and the distribution of pRFs sparser in the peripheral visual field, as qualified by the saturation weighting in our searchlight back-projections. This might have additionally contributed to the unclear patterns in V1 and V2. Nevertheless, our validation analyses showed that when contrasting the vertical or horizontal condition to fixation, we were able to effectively stimulate the cortical area corresponding to the top-right visual field quadrant. This confirms the general feasibility of our back-projection approach.

## 6. General discussion

In three experiments, we used dynamic bistable (diamond experiment) and non-ambiguous stimuli (dots and dots quadrant experiment) to explore within- and between-area response amplitude mechanisms underlying global object perception in human visual cortex. All these stimuli could either be perceived globally (i.e., as a grouped and coherently moving Gestalt) or locally (i.e., as ungrouped and incoherently moving elements).

### 6.1. Signatures in lower visual cortex

When contrasting global to local perception, our diamond and dots experiment revealed a fairly wide-spread suppression of activity across the whole visual field in lower visual cortex. However, unlike our diamond experiment, our dots experiment provided little evidence for pronounced activity modulations in the background region, suggesting that these suppressive effects might be diffusely related to the physical stimulus. Our dots quadrant experiment largely confirmed this notion, but revealed additionally a wide-spread increase of activity in the background area. Whereas the wide-spread suppressive effects from the diamond experiment speak against a within-area response amplitude mechanism mediating global object perception, the results from the dots and dots quadrant experiment are largely compatible with this idea. In any case, the outcomes of our experiments seem to converge in that they suggest that perceptual grouping results in a reduction of activity in lower visual cortex.

Surprisingly, however, all these findings are at odds with recent evidence showing a decrease of brain activity in the background and stimulus region of another bistable global-local stimulus along with an increase in the center and inferred contour region for global vs local perception (Grassi et al., 2017). Unlike our diamond stimulus, this bistable stimulus triggers a local percept of four individually rotating disk pairs or a global percept of two floating squares circling around the stimulus center. The mismatch in findings might therefore be related to differences in physical stimulus properties, such as the type and/or direction of motion (i.e., rotary vs oscillatory and rotational vs horizontal/vertical, respectively).

The emergence of suppressive effects in the dots and dots quadrant experiment, where shape information was kept constant during global and local perception, further highlights the importance of motion properties. This idea is in line with findings of reduced activity in lower visual cortex for coherent vs incoherent motion (Braddick et al., 2001; Costagli et al., 2014; Harrison et al., 2007; McKeefry et al., 1997; Schindler & Bartels, 2017), although no or opposite effects have also occasionally been observed (Braddick et al., 2001; Rees et al., 2000). However, unlike these studies on motion coherence, we did not compare coherent to random motion nor did Grassi et al. (2017). Rather, all our stimuli always comprised coherent motion, but were either perceived as ungrouped and moving out-of-phase (local) or grouped and moving in-phase (global). Accordingly, although speculative, the perceived axis of motion (horizontal vs vertical) might constitute an important factor driving our results.

A potential reason for a horizontal-vertical imbalance might be that there is a bias for vertical motion in lower visual cortex resulting in generally higher response amplitudes. In the case of the diamond experiment (in particular), this directional anisotropy might additionally interact with feature-based attention. Specifically, given that information about motion direction is inherently ambiguous for the diamond stimulus, during the local diamond state, observers may direct their attention to vertical motion and during the global diamond state to horizontal motion.

Interestingly, there is evidence for increased responses to horizontal/vertical motion around the horizontal/vertical meridian in lower visual cortex (Clifford et al., 2009). Along with a plethora of similar studies (Maloney et al., 2014; Raemaekers et al., 2009; Schellekens et al., 2013), this finding points to a radial response bias. Importantly, such a radial anisotropy is incompatible with our results, as it would produce meridian-related antagonistic effects for global as compared to local perception (i.e., an increase in differential activity around the horizontal meridian and decrease around the vertical meridian), which we did not observe. Critically, however, it is hitherto not clear in how far these radial anisotropies are due to vignetting (Roth et al., 2018) and/or aperture-inward biases (Wang et al., 2014), leaving open the possibility for a vertical-horizontal anisotropy.

The role of feature-based attention as a perceptual modulator fits in with evidence that the attended direction of motion can be decoded from activity in lower visual cortex (Kamitani & Tong, 2006) even in the absence of direct physical stimulation (Serences & Boynton, 2007) and the idea that feature-based attention acts fairly globally across the visual field (Jehee et al., 2011; Maunsell & Treue, 2006; Saenz et al., 2002; Serences & Boynton, 2007; Treue & Martinez Trujillo, 1999). Strikingly, the combinatory effect of anisotropies and feature-based attention might also help explain why variations of the diamond stimulus triggering a local percept of vertical motion and a global percept of rotational motion (Caclin et al., 2012) or other bistable global-local stimuli (Grassi et al., 2017) produce distinct differential response profiles. Most importantly, as for our findings, this combinatory effect leads to the prediction that rotating the diamond display by 90 degree should produce the opposite pattern of results for global vs local perception.

Leaving all inconsistencies aside, our study overlaps with studies on motion coherence (Braddick et al., 2001; Costagli et al., 2014; Harrison et al., 2007; McKeefry et al., 1997; Schindler & Bartels, 2017) and Grassi et al.’s (2017) work in that it points to *stimulus-referred* suppressive effects for global vs local perception. This suppression might be related to a recently reported phenomenon known as the *global slow-down effect* (Kohler et al., 2009, 2014). This effect comprises a slow-down in the perceived speed of a stimulus configuration as a result of perceptual grouping and has hitherto only been demonstrated behaviourally (Kohler et al., 2009, 2014) for variations of the stimulus used by Grassi et al. (2017). As such, it would be worthwhile to examine whether the effect holds true for the diamond stimulus and ultimately also our dots and dots quadrant stimuli along with more conventional motion displays because these stimulus classes abstract from shape perception (for a similar point and a discussion on potential underlying mechanisms see Kohler et al., 2014).

The broad background enhancement we observed in the dots quadrant experiment, which was absent in the diamond and dots experiment, might be due to spatial attention. In particular, perceiving a grouped and coherently moving object parafoveally might require fewer attentional resources than perceiving an ungrouped and incoherently moving object. Accordingly, in the vertical condition, fewer attentional resources might have been available for processing the background area. This interpretation fits in with reports that spatial attention results in increased brain responses even in the absence of physical stimulation (Kastner et al., 1999; Silver et al., 2009). Due to the size and central presentation of the diamond and dots stimulus, we might have been unable to observe similar effects in the diamond and dots experiment. It is furthermore possible that the background enhancement is related to *perceived back-ground luminance*, which has recently been found to be increased for global vs local perception (Han & VanRullen, 2016, 2017).

Building upon previous research involving the diamond stimulus (De-Wit et al., 2012), it is important to highlight that our results in lower visual cortex across all experiments contradict suggestions of predictive coding theories that suppressive effects should be confined to cortical sites encoding the physical stimulus and accompanied by unchanged activity in the background region (e.g., Mumford, 1992; Murray et al., 2004; Rao & Ballard, 1999). They furthermore conflict with alternative accounts, such as *response sharpening* (e.g., Kersten et al., 2004; Kersten & Yuille, 2003; Murray et al., 2004). Response sharpening accounts assume that predictive feedback from higher-tier areas sharpens diffuse responses in lower-tier areas (due to noise or ambiguity) by increasing activity matching the global interpretation of the bottom-up input and decreasing non-matching activity. Accordingly, when contrasting global to local object perception, activity should increase in stimulated and decrease in non-stimulated sites – a pattern we did not observe.

### 6.2. Relationship between higher and lower visual cortex

Whereas our findings for the VLOC in the dots and dots quadrant experiment largely paralleled those in lower visual cortex for global vs local perception, we observed a large-scale response enhancement in the diamond experiment that was antagonistic to responses in lower visual cortex. The absence of an inverse relationship between lower visual cortex and the VLOC when shape information did not change suggests that this between-area response amplitude code does not represent a generic grouping mechanism acting beyond shape perception.

It could be argued that our failure to find evidence for such an opposite pattern is due to the fact that non-ambiguous stimuli strongly favor a single perceptual interpretation and thus involve less predictive feedback (Wang et al., 2013). This explanation seems unlikely because an inverse V1-LOC relationship has also been established for non-ambiguous shape-like stimuli vs unstructured displays (Murray et al., 2002). Moreover, at least broadly in line with our results, recent studies (Grassi et al., 2016, 2018) found no evidence for the involvement of the LOC when a dynamic, bistable global-local stimulus constantly triggered shape-based interpretations (i.e., moving disks forming large squares or small circles).

The absence of a (stimulus-related) increase in VLOC activity in the dots and dots quadrant experiment seems incompatible with a study reporting enhanced LOC activity for intact compared to scattered objects with disturbed inter-part relations (Margalit et al., 2017). Yet, in this study, inter-part relations were abolished by disturbing the contiguity of different shape parts. In our experiments, however, the position of the apertures did not change during the local state nor did shape information, which might explain the discrepant results.

### 6.3. Inter-individual variability

The wealth of evidence presented here is based on data pooled across a small number of participants. As such, it is important to flag an overly large influence of a single participant. Although the results of our representational similarity analyses did not indicate such a bias, they collectively highlighted the idiosyncratic nature of the individual back-projection profiles. Some of these idiosyncrasies are likely due to a lower signal-to-noise ratio at the individual level triggered by a generally lower number of available data points. They might also be related to inter-individual variability in pRF estimates and processing of the global-local stimuli, such as differences in switch rates, perceptual durations (Supplementary material, 1.1.2 Results, and Supplementary Figure S2), perceptual vividness, and attention allocation.

## 7. Conclusion

We found evidence for a suppression of activity in lower visual cortex accompanied by an increase of activity in the VLOC for global relative to local object perception. While the suppressive effects in lower visual cortex manifested themselves irrespective of shape grouping, this was not the case for the enhanced responses in the VLOC. Instead, once shape perception was held constant during both global and local object perception, the VLOC also showed a decrease of activity. As such, the inverse relationship between lower visual cortex and the VLOC we initially quantified cannot be regarded as a generic grouping mechanism. We furthermore observed that grouping-related suppressive effects can be diffusely confined to stimulated visual field portions (once stimulus size is reduced) and surrounded by enhancement effects, potentially pointing to a within-area response amplitude mechanism mediating the perception of figure and ground.

## 8. Data and code availability

Preprocessed data, analysis code, and stimulus videos are available from https://doi.org/10.17605/OSF.IO/E6C8S.

## 10. Acknowledgements

This research was supported by a European Research Council Starting Grant to DSS (WMOSPOTWU, 310829). We thank Elisa Infanti and Man-Ling Ho for their help with data collection, Samuel G. Solomon and Matteo Lisi for useful discussions, and all participants for taking part in our study. Some of the reports here were part of an unpublished MSc thesis.

## 1. Supplementary methods and results

### 1.1. Diamond experiment

#### 1.1.1. Data analysis

##### Perceptual durations

Participants’ key presses were used to calculate the durations of the diamond and no-diamond percept. If the same key was pressed multiple times in succession, the resulting sub-durations were summed up. The period from the onset of the diamond display until participants’ first key press was discarded. For each participant and the data pooled across participants, we then fit the durations for the diamond and no-diamond percept with a two-parameter (α: shape, β: rate) gamma probability density function using the maximum likelihood method. The resulting fits were superimposed onto a density histogram of the perceptual durations (bin width: 2 s).

#### 1.1.2. Results

##### Perceptual durations

The probability density histograms of the durations per perceptual state for each participant and the pooled data with superimposed gamma fit can be found in Supplementary Figure S2. Despite inter-individual variability in the shape and rate parameters, both the pooled and individual diamond and no-diamond durations seem to be well fit with a gamma distribution, suggesting they follow similar temporal dynamics. However, all participants except P2 showed a tendentially higher probability density of longer durations for the no-diamond relative to the diamond percept. Likewise, these participants showed a higher median duration for the no-diamond percept and spent a higher proportion of time in this perceptual state, which was also reflected in the pooled results. Consequently, the perception of most participants was slightly biased towards the no-diamond state.

## Supplementary figures and tables

**Figure S1.**
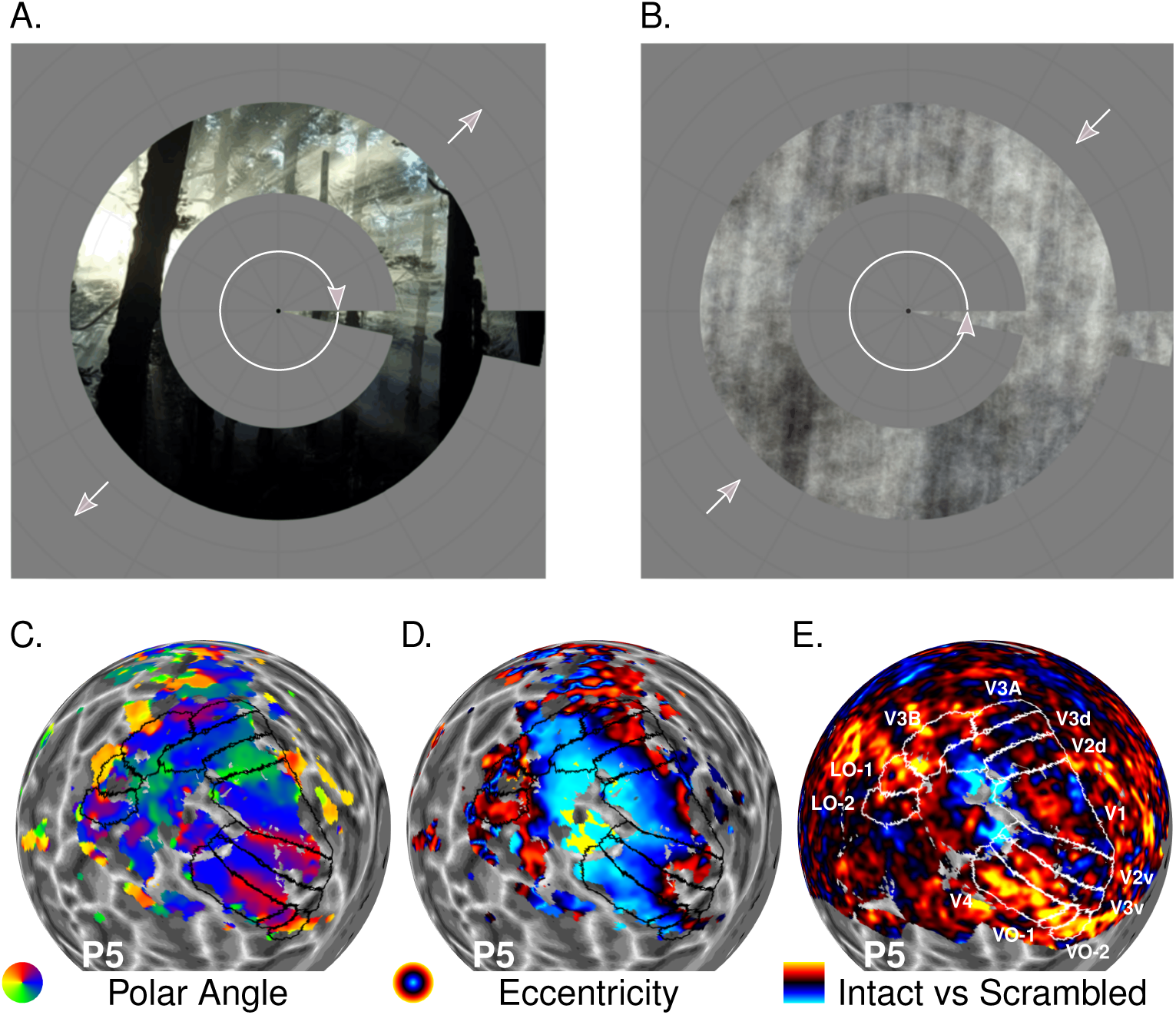
Retinotopic mapping experiment. Example frames of the wedge-and-ring stimulus and smooth cortical maps of P5’s left hemisphere projected onto a spherical surface model. **A.** Intact, colorful carrier pattern. **B.** Phase-scrambled version of the carrier pattern in A. The gray arrows indicate the movement direction of the wedge-and-ring aperture, which was either clockwise and expanding (A.), counterclockwise and contracting (B.), clockwise and contracting, or counterclockwise and expanding (not shown, respectively). **C.** Polar angle map. **D.** Eccentricity map. Vertices surpassing an eccentricity of 15 dva were discarded (no other post-smoothing thresholding was applied). Note that these pRF maps were subjected to the experiment-specific smoothing procedure (see 3.1.7 Data analysis). The color disks represent the color schemes used to label different visual field portions. **E.** Differential brain activity resulting from contrasting periods of intact vs phase-scrambled images. Differential betas surpassing a value of ± 2 were set to that value. Cold colors reflect negative and warm colors positive differential beta values as indicated by the color bar. White or black lines denote the boundaries between visual areas. The gray scale pattern of the surface model reflects the cortical curvature. Darker regions depict sulci and lighter regions gyri. P5 = Participant 5. VO = Ventral-occipital area. LO = Lateral-occipital area. PRF = Population receptive field.

**Figure S2.**
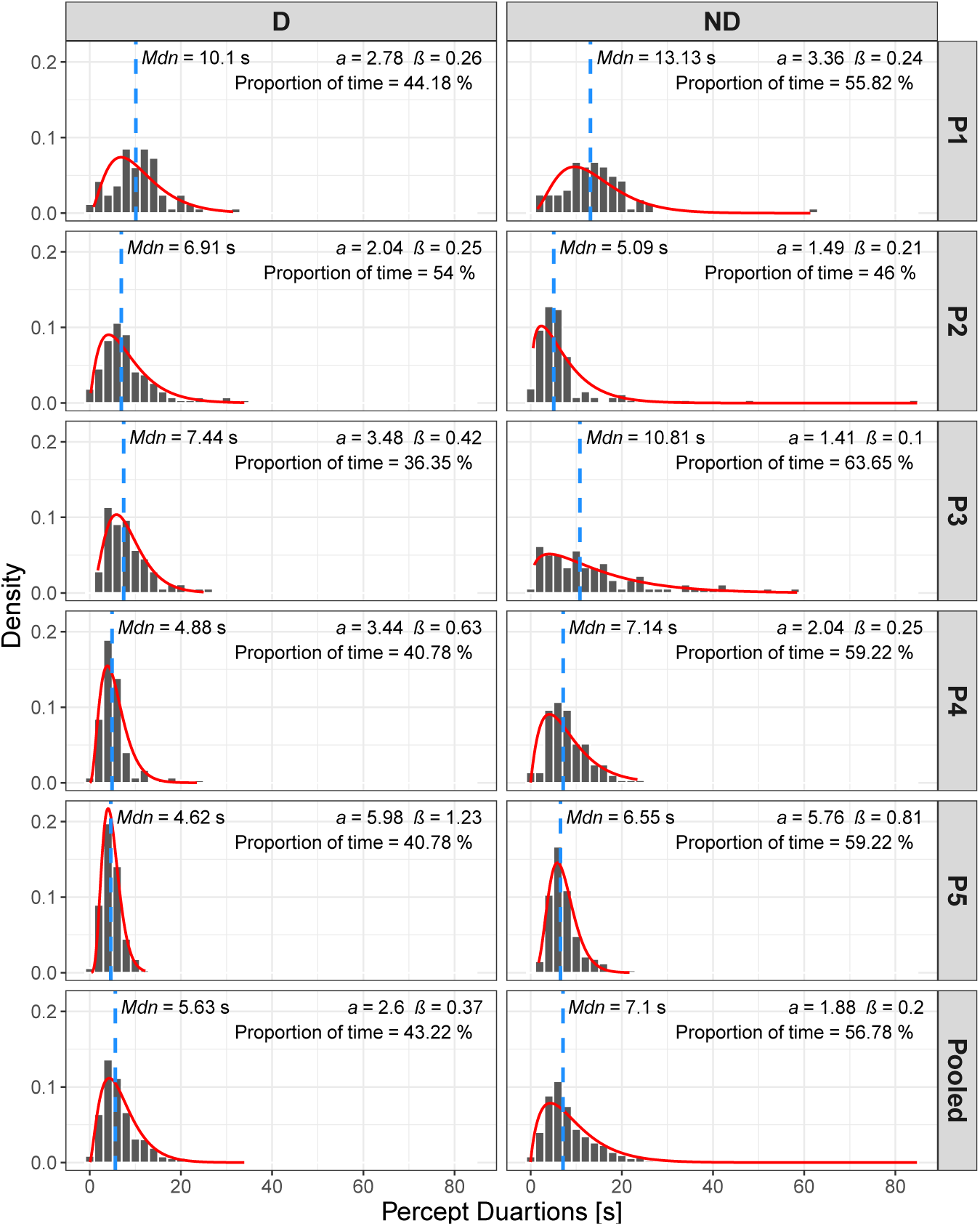
Diamond experiment. Probability density histograms of the durations corresponding to the diamond and no-diamond percept with superimposed gamma functions. The red line depicts the fitted gamma curve and the blue line the median duration. *α*, *β* = Shape and rate parameter of the gamma distribution, respectively. Total time = Proportion of time spent in the respective perceptual state. D = Global, diamond percept. ND = Local, no-diamond percept. P1-P5 = Participant 1-5. Pooled = Data pooled across all 5 participants.

**Figure S3.**
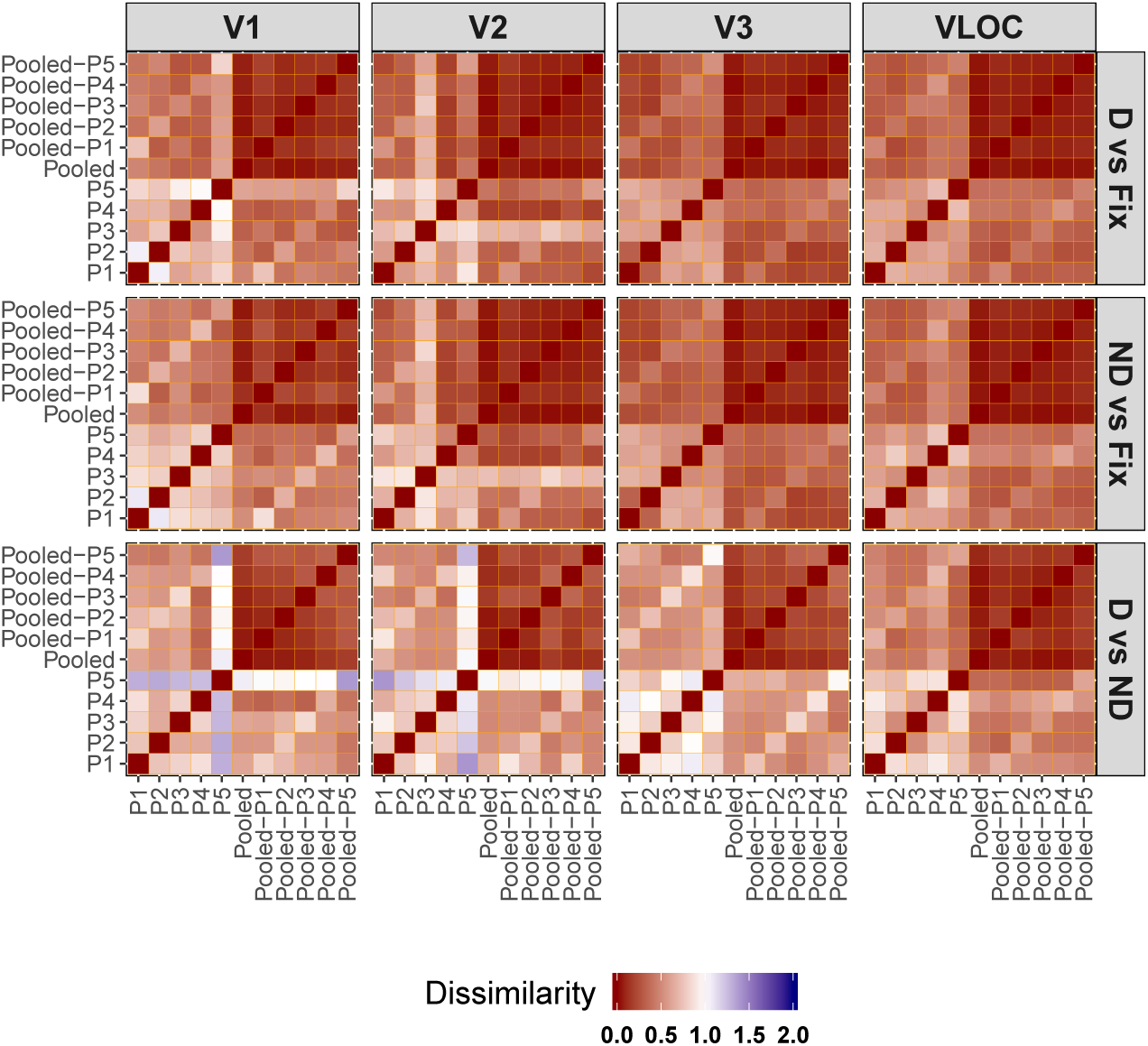
Diamond experiment. Representational dissimilarity matrices for the individual, pooled, and LOSO searchlight back-projections as a function of contrast of interest and visual area. Dissimilarities were defined as 1-Spearman correlation. D = Global, diamond percept. ND = Local, no-diamond percept. Fix = Fixation baseline. VLOC = Ventral-and-lateral occipital complex. P1-P5 = Participant 1-5. Pooled = Data pooled across all 5 participants. Pooled-P1-Pooled-P5 = Data pooled across 4 participants with 1 participant left out (as indicated by the suffix). LOSO = Leave-one-subject-out.

**Figure S4.**
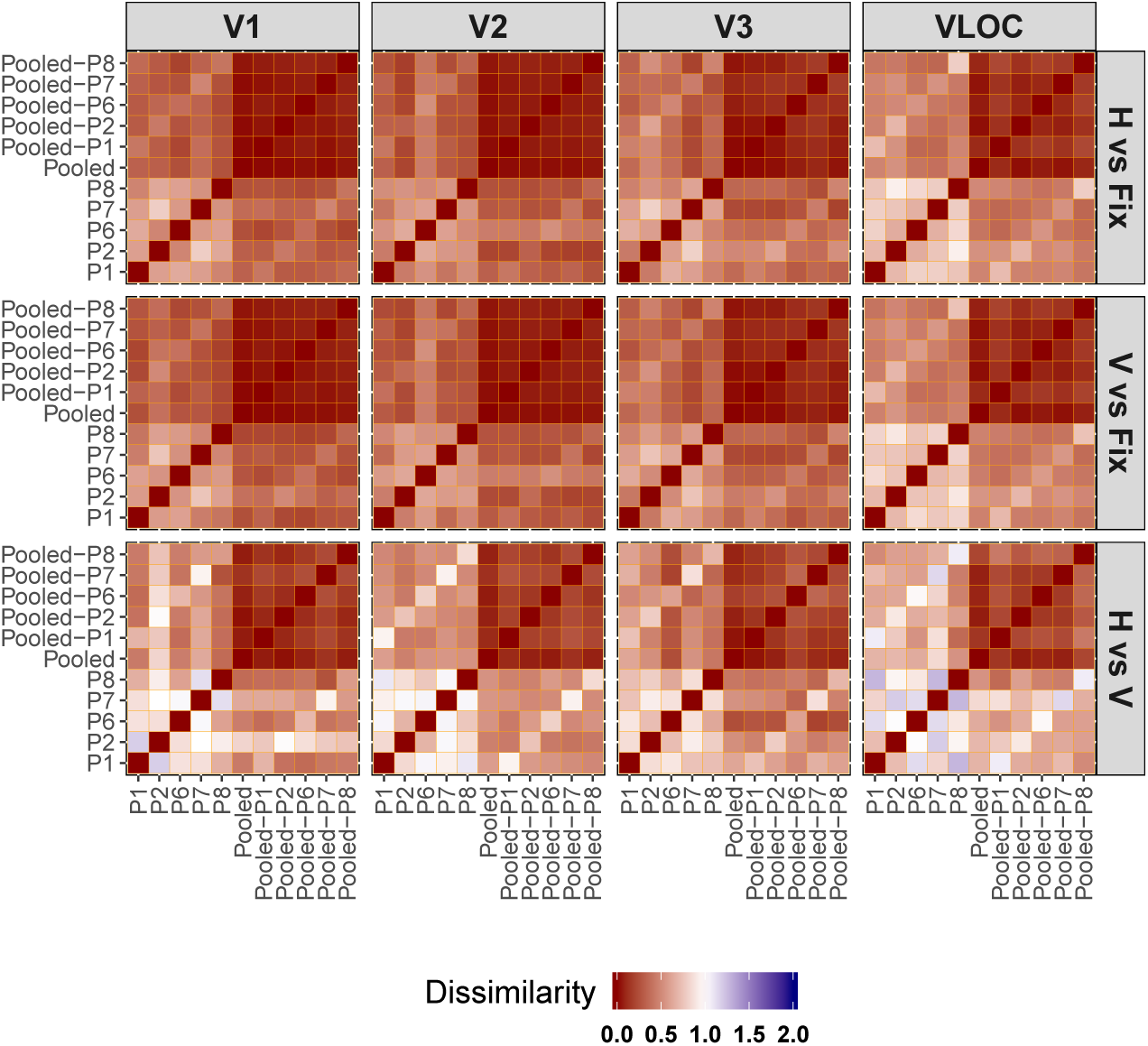
Dots experiment. Representational dissimilarity matrices for the individual, pooled, and LOSO searchlight back-projections as a function of contrast of interest and visual area. Dissimilarities were defined as 1-Spearman correlation. H = Global, horizontal condition. V = Local, vertical condition. Fix = Fixation baseline. VLOC = Ventral-and-lateral occipital complex. P1-P2 and P6-P8 = Participant 1-2 and 6-8. Pooled = Data pooled across all 5 participants. Pooled-P1-Pooled-P2 and Pooled-P6-Pooled-P8 = Data pooled across 4 participants with 1 participant left out (as indicated by the suffix). LOSO = Leave-one-subject-out.

**Figure S5.**
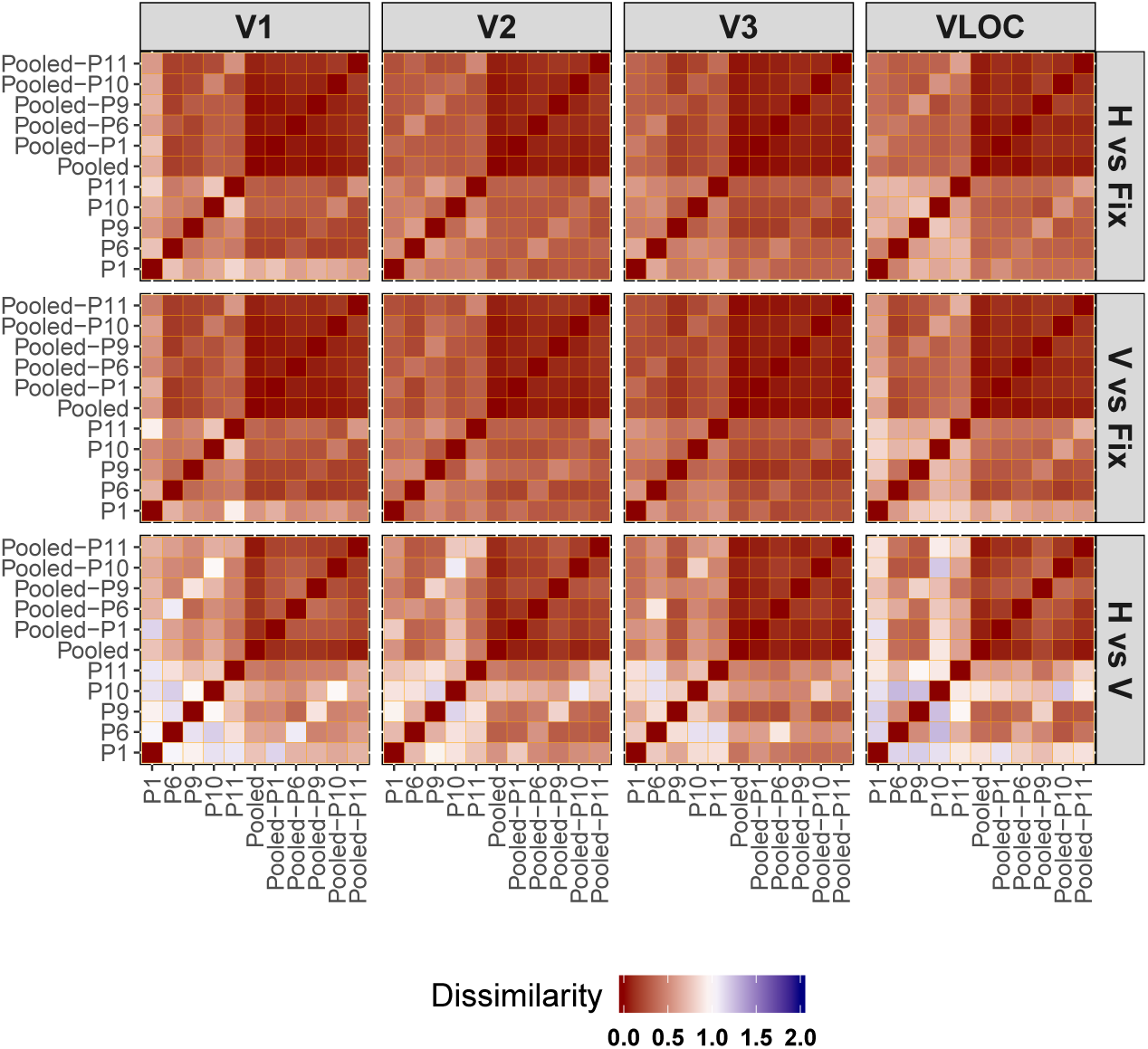
Dots quadrant experiment. Representational dissimilarity matrices for the individual, pooled, and LOSO searchlight back-projections as a function of contrast of interest and visual area. Dissimilarities were defined as 1-Spearman correlation. H = Global, horizontal condition. V = Local, vertical condition. Fix = Fixation baseline. VLOC = Ventral-and-lateral occipital complex. P1, P6, P9-P11 = Participant 1, 6, and 9-11. Pooled = Data pooled across all 5 participants. Pooled-P1, Pooled-P6, Pooled-P9-Pooled-P11 = Data pooled across 4 participants with 1 participant left out (as indicated by the suffix). LOSO = Leave-one-subject-out.

**Table S1.** Software and Toolboxes

| Implemented procedures | Software (version) | Toolbox (version) |
| --- | --- | --- |
| Experiments | Matlab R2014a (8.3)<sup>1</sup> | PTB (3.0.11)<sup>4</sup> |
| Preprocessing |  |  |
| Realignment/unwarping/coregistration | Matlab R2016b (9.1)<sup>1</sup> | SPM8 (6313; default parameters)<sup>5</sup> |
| Surface reconstruction | FreeSurfer (5.3.0)<sup>2</sup> |  |
| Surface projection/detrending/standardization | Matlab R2016b (9.1)<sup>1</sup> | SamSrf (5.84)<sup>6</sup> |
| pRF estimation | Matlab R2016b (9.1)<sup>1</sup> | SamSrf (5.84)<sup>6</sup> |
| Delineations | Matlab R2016b (9.1)<sup>1</sup> |  |
| GLM |  | SPM8 (6313)<sup>5</sup> |
| Surface projection/manual demarcation |  | SamSrf (5.84)<sup>6</sup> |
| Smoothing/visualizations |  | SamSrf (6.20)<sup>7</sup> |
| Searchlight back-projections | Matlab R2016b (9.1)<sup>1</sup> |  |
| GLM |  | SPM8 (6313)<sup>5</sup> |
| Surface projection |  | SamSrf (5.84)<sup>6</sup> |
| Smoothing/searchlight algorithm/visualizations |  | SamSrf (6.20)<sup>7</sup> |
| Representational similarity |  |  |
| Dissimilarity calculation | Matlab R2016b (9.1)<sup>1</sup> |  |
| NMDS/visualizations | R (3.5.3)<sup>3</sup> | vegan (2.5-6)<sup>8</sup><br>ggplot2(3.2.1)<sup>9</sup><br>reshape2(1.4.3)<sup>10</sup><br>plyr (1.8.4)<sup>11</sup><br>rmatio (0.14.0)<sup>12</sup> |
| Perceptual durations |  |  |
| Duration calculation | Matlab R2016b (9.1)<sup>1</sup> |  |
| Gamma fitting/visualizations | R (3.5.3)<sup>3</sup> | MASS (7.3-51.4)<sup>13</sup><br>ggplot2 (3.2.1)<sup>9</sup><br>rmatio (0.14.0)<sup>12</sup> |

Note.<sup>1</sup>https://uk.mathworks.com/. <sup>2</sup>Dale et al. (1999) and Mendola et al. (1999). <sup>3</sup>R Core Team (2018). <sup>4</sup>Brainard (1997), Kleiner et al. (2007), and Pelli (1997). <sup>5</sup>https://www.fil.ion.ucl.ac.uk/spm/software/spm8/. <sup>6</sup>https://doi.org/10.6084/m9.figshare.1344765.v24. <sup>7</sup>https://osf.io/s3h7w/. <sup>8</sup>Oksanen et al. (2019). <sup>9</sup>Wickham (2016). <sup>10</sup>Wickham (2007). <sup>11</sup>Wickham (2011). <sup>12</sup>Widgren & Hulbert (2019). <sup>13</sup>Kafadar et al. (1999). pRF = population receptive field. GLM = General linear model. NMDS = Non-metric multidimensional scaling. PTB = Psychtoolbox.

